# SLOW5: a new file format enables massive acceleration of nanopore sequencing data analysis

**DOI:** 10.1101/2021.06.29.450255

**Authors:** Hasindu Gamaarachchi, Hiruna Samarakoon, Sasha P. Jenner, James M. Ferguson, Timothy G. Amos, Jillian M. Hammond, Hassaan Saadat, Martin A. Smith, Sri Parameswaran, Ira W. Deveson

## Abstract

Nanopore sequencing is an emerging genomic technology with great potential. However, the storage and analysis of nanopore sequencing data have become major bottlenecks preventing more widespread adoption in research and clinical genomics. Here, we elucidate an inherent limitation in the file format used to store raw nanopore data – known as FAST5 – that prevents efficient analysis on high-performance computing (HPC) systems. To overcome this, we have developed SLOW5, an alternative file format that permits efficient parallelisation and, thereby, acceleration of nanopore data analysis. For example, we show that using SLOW5 format, instead of FAST5, reduces the time and cost of genome-wide DNA methylation profiling by an order of magnitude on common HPC systems, and delivers consistent improvements on a wide range of different architectures. With a simple, accessible file structure and a ^~^25% reduction in size compared to FAST5, SLOW5 format will deliver substantial benefits to all areas of the nanopore community.

## MAIN TEXT

The emergence of nanopore sequencing is reshaping the landscape of genomics. Devices from Oxford Nanopore Technologies (ONT) enable sequencing of native DNA and RNA molecules with no theoretical upper limit on read length^1^. This supports the accurate assembly and phasing of repetitive genomes and metagenomes^2–6^; enhanced detection of structural and/or complex genetic variation^7–11^; assembly-free detection and quantification of spliced RNA transcripts^12^; and profiling of diverse epigenetic and RNA modifications^13–18^. High-output benchtop ONT instruments (GridION and PromethION) have recently enabled cost-effective nanopore sequencing of human and other large eukaryotic genomes^7,8,19^. Yet, with sequencing throughput rapidly increasing, large data volumes and computational bottlenecks have become a major impediment.

ONT devices measure the displacement of ionic current as a DNA or RNA strand passes through a biological nanopore, recording time-series signal data in FAST5 format (**Figure 1a**; **Supplementary Note 1**). This signal data is typically translated, or ‘base-called’, into sequence reads (FASTQ or FASTA format) prior to downstream analysis. Many bioinformatics tools also directly access the signal data in order to improve the accuracy of assembled genomes or detect fine signal-perturbations that are indicative of DNA/RNA modifications, genetic variants or other features (**Figure 1a**)^5,14,16–18^. However, nanopore signal data is very large (^~^1.3 TB FAST5 files for a ^~^30X human genome; **Supplementary Table 1**) and both base-calling and downstream analysis steps involving FAST5 files are computationally expensive.

**Figure 1.**
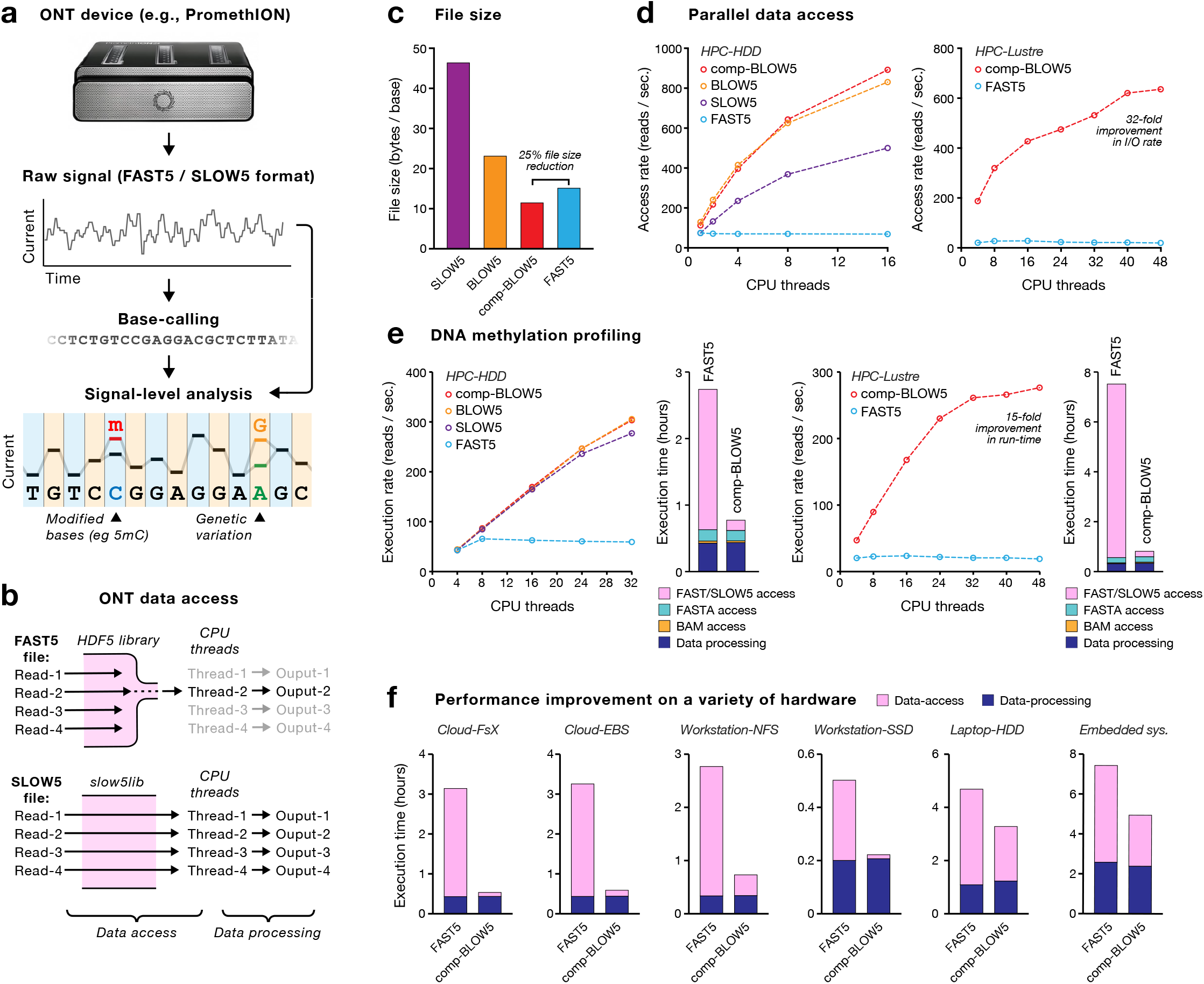
SLOW5 format enables efficient parallel analysis of nanopore signal data. (**a**) Schematic illustrates the typical lifecycle of nanopore data. Raw current-signal data is generated on an Oxford Nanopore Technologies (ONT) sequencing device and written in FAST5 format. Raw data is base-called into sequence reads (FASTQ/FASTA format). Downstream analysis involving both base-called reads and raw signal data is used to identify genetic variants, epigenetic modifications (e.g., *5mC*) and other features. (**b**) Schematic illustrates the bottleneck in ONT signal data analysis. FAST5 file reading requires the HDF5 software library, which serialises file-access requests by multiple CPU threads, preventing efficient parallel analysis. SLOW5 files are not dependent on the HDF5 library and are amenable to efficient parallel analysis. A more detailed mechanistic diagram is provided in **Fig. S1e**. (**c**) Bar chart shows the relative file sizes of ASCII SLOW5 (purple), binary BLOW5 (orange) and compressed-BLOW5 (comp-BLOW5; red) formats, compared to FAST5 (blue). (**d**) Dot plots show the rate of file access (reads / second) for the above file types, as a function of CPU threads used on two HPC systems: *HPC-HDD* (left) or *HPC-Lustre* (right). (**e**) Dot plots show the rate of execution (reads / second) for DNA methylation calling for the same file types on *HPC-HDD* (left) and *HPC-Lustre* (right). For the instance of maximum CPU threads, bar charts show the time consumed by individual workflow components: FAST5/SLOW5 data access (pink), FASTA data access (teal), BAM data access (orange) and data processing (navy). (**f**) Bar charts show the time consumed by data access (pink) and data processing (navy) during DNA methylation calling on a range of different computer systems. Full specifications are provided in **Supplementary Table 2.**

Currently, the most popular signal-level analysis among nanopore users is DNA methylation profiling with the software *Nanopolish*^17^. We therefore selected this example use-case as the basis for a forensic investigation of FAST5 data analysis on typical high-performance computing (HPC) systems (**Supplementary Note 2**). FAST5 is a Hierarchical Data Format 5 (HDF5) file with a specific schema defined by ONT. HDF5 is a complex file format for storing large data that can only be read and written using a single [software library first developed in 1998](https://support.hdfgroup.org/about/history.html). Our investigations revealed that: (*i*) the use of increasing numbers of parallel CPU threads resulted in a relatively small reduction in the overall run-time of a typical *Nanopolish* job (**Fig. S1a**); (*ii*) this was due to inefficient data access (file reading), rather than inefficient data processing (**Fig. S1a-d**); (*iii*) the underlying bottleneck was a limitation in the software library for reading HDF5 files that prevents efficient utilisation of parallel CPU resources (**Fig. S1e**; **Supplementary Note 2**).

Parallel computing enables scalable analysis of large datasets and is central to modern genomics. Regrettably, our investigation shows that FAST5 format suffers from an inherent inefficiency that ensures, even with access to the most advanced HPC systems, analysis of nanopore signal data will be prohibitively slow (**Figure 1b).** For example, with the maximum resource allocation available on Australia’s National Computing Infrastructure (among the world’s largest academic supercomputers; see **Supplementary Table 2**; HPC-Lustre), genome-wide DNA methylation profiling on a single ^~^30X human genome sequencing dataset runs for >14 days at a cost of >$500. Moreover, given that the vast majority (>90%) of the overall run-time is simply spent reading FAST5 files, further optimization to the *Nanopolish* software (or any other software that reads FAST5 files) would have minimal benefit.

To overcome the inherent limitations in FAST5 format, we created SLOW5; a new file format that is designed for efficient, scalable analysis of nanopore signal data (**Figure 1b**). SLOW5 format encodes all information found in FAST5 format but is not dependent on the HDF5 library required to read FAST5 files. The human readable version of SLOW5 format is a tab-separated values (TSV) file encoding metadata and time-series signal data for one ONT read per line, with global metadata stored in a file header (**Table 1**; **Supplementary Note 3**). Parallel file access is facilitated by an accompanying binary index file that specifies the position of each read (in bytes) within the main SLOW5 file (**Supplementary Note 3**). SLOW5 can be encoded in human-readable ASCII format, or a more compact and efficient binary format, BLOW5; analogous to the seminal SAM/BAM format for storing DNA sequence alignments^20^. The binary format optionally supports zlib compression (compressed-BLOW5), thereby minimising the data storage footprint while still permitting efficient parallel access. Compressed-BLOW5 format is ^~^25% smaller in size than FAST5 format (11.63 vs 15.30 bytes/base; **Figure 1c**). For a single ^~^30X human genome dataset, this equates to a reduction of ^~^300 Gbytes and a saving ^~^$50 per month on Amazon cloud storage. We also note that substantial further space savings can likely be achieved with alternative compression methods (e.g., [zstd](https://github.com/facebook/zstd)).

**Table 1:**
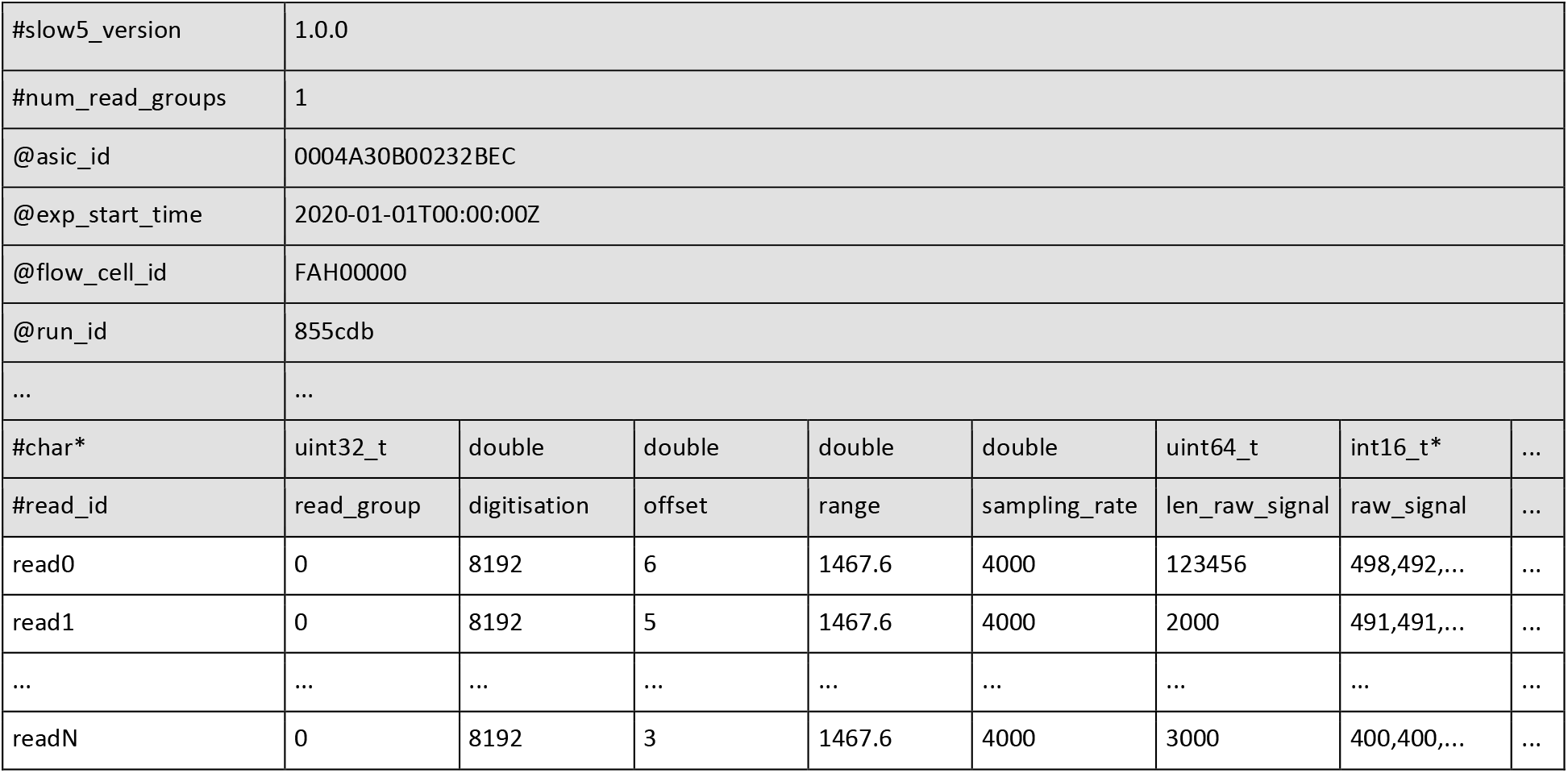
Example of a SLOW5 ASCII file with a single read group. A SLOW5 file contains a header (grey rows; ‘@’ and ‘#’ prefixes) that stores metadata regarding the contents of the file and the ONT experiment/s contained within, followed by data records (white rows; no prefixes) for sequencing reads, with one read per line. In this example, column/row borders are added to increase the readability. The actual format uses tabs (‘\t’) and newlines (‘\n’) as delimiters. Complete format specifications are provided in **Supplementary Note 3**.

To determine the performance benefits of SLOW5 format, we first measured data access using a small human DNA sequencing dataset of ^~^500 million reads (**Supplementary Table 1**) on two different HPC systems (HPC-HDD and HPC-Lustre; **Supplementary Table 2**). The rate of SLOW5 data access (reads/second) was faster than FAST5 across the board, and increased with the use of additional CPU threads, whereas FAST5 access was largely unchanged (**Figure 1d**). This trend, which reflects the capacity of SLOW5 (but not FAST5) to be efficiently accessed by multiple CPU threads in parallel, was observed regardless of whether data was stored in SLOW5, BLOW5 or compressed-BLOW5 format, with the latter exhibiting the most efficient data access (**Figure 1d**). As a result, we observed substantial improvements in data access rates when using many CPUs on both HPC systems. For example, using 48 CPU threads on the HPC-Lustre system, ^~^20 hours were required to read this small dataset in FAST5 format, compared to just ^~^13 minutes in compressed-BLOW5 (^~^32-fold improvement; **Figure 1d**).

This improvement in the efficiency of data access manifested in substantial performance gains during DNA methylation calling. When using SLOW5 input data, the time taken to run *Nanopolish* was reduced in proportion to the number of CPUs available on both HPC systems (**Figure 1e**). This is indicative of efficient parallel computation and was not observed when using FAST5 (**Figure 1e**). As a result, substantial performance improvements were observed when using many CPUs, with a maximum ^~^15-fold reduction in run-time with 48 CPUs on the HPC-Lustre system (**Figure 1e**). The improvement is the result of efficient data-access, with no difference observed in data-processing between the different file formats (**Fig. S2a,b**). Indeed, whereas data-access was the major bottleneck for FAST5 analysis, it constituted a negligible fraction of the total run-time for SLOW5 analysis (**Fig. S2c,d**). Put simply, this means that overall performance is dictated by the efficiency of the program, rather than simply the time taken to read the input data, thereby enabling optimisation through further engineering. For example, using a GPU-accelerated version of *Nanopolish* (described elsewhere^21^), with compressed-BLOW5 input data, we were able to complete whole-genome methylation profiling on a single 30X human dataset in just ^~^10.5 hours with 48 threads (>30-fold improvement compared to standard analysis with FAST5 input; **Supplementary Table 2**; HPC-GPU).

While SLOW5 format is designed to enable scalable analysis on multi-CPU HPC systems, we reasoned that improved data access would be beneficial on almost any computer. To test this, we benchmarked DNA methylation profiling, as above, on a wide range of different architectures (**Supplementary Table 2**). In all cases, the time consumed by data access was significantly reduced, leading to overall improvements in execution time (**Figure 1f**). As expected, performance improvements were greatest on systems with larger numbers of CPUs, such as a cloud-based virtual machine on Amazon AWS (^~^7-fold improvement at 32 CPU threads). However, meaningful benefits were observed even on miniature devices for portable computing, such as an NVIDIA Xavier embedded module (^~^60% improvement; **Figure 1f**). In summary, the use of SLOW5 delivered substantial improvements during DNA methylation profiling on a diverse range of hardware.

To ensure FAST5 to SLOW5 file conversion is not a barrier to SLOW5 adoption (given ONT devices write data in FAST5 by default), we implemented software for efficient, parallelisable, lossless conversion from FAST5 to SLOW5, as well as conversion between different SLOW5 formats. File conversion times are proportionally reduced with high CPU availability, and are trivial compared to execution times for typical FAST5 analysis (**Fig. S3a,b**). For example, conversion of a single ^~^30X human genome dataset from FAST5 to compressed-BLOW5 takes just ^~^3 hours with 48 CPUs. We additionally implemented software for live FAST5 to SLOW5 file conversion during a sequencing run, utilising the internal computer on an ONT PromethION device (**Fig. S3c**). This means the user can obtain raw data from their ONT experiment in compressed-BLOW5 format with effectively zero additional workflow hours required for file conversion.

The inefficiency of data access on FAST5 files creates substantial delays and expenses, limiting the feasibility of ONT sequencing for many applications in research and clinical genomics. Arguably, these frictions have also acted to discourage development of bioinformatics software that directly accesses nanopore signal data. Given the complexity and cost of working with FAST5 files, active work in this area has been limited to a small number of well-resourced groups. This is in stark contrast to the simple, efficient and open-source SAM/BAM sequence alignment format, developed in 2009^20^, that was a key catalyst in the growth of genome informatics and remains the central file format.

SLOW5 format provides the framework for efficient, parrelisable analysis of nanopore signal data for any intended application. SLOW5 reading/writing is managed by efficient software APIs for both the C (*slow5lib*) and python (*pyslow5*) languages. This facilitates the integration of SLOW5 into third-party bioinformatics software, including integration into existing packages by simply replacing the existing API for FAST5 reading/writing. The SLOW5 API is fully open source and not dependent on the archaic HDF5 library required to read FAST5 files. Along with the simple, intuitive structure of SLOW5 format, this will support active and open software development for ONT data analysis.

## METHODS

### Reading and writing SLOW5 files: slow5lib

*slow5lib* is implemented using the C programming language. To maximise portability, the *slow5lib* code follows the C99 standard with X/Open 7 POSIX 2008 extensions. Sequential access to SLOW5 ASCII files and SLOW5 binary files is performed using the *getline()* and *fread()* functions respectively. For performing random disk accesses to SLOW5, the SLOW5 index is first loaded to a hash table in RAM. The read-identifier serves as the hash table key. For a given read-identifier, the file offset and the record length are obtained from this hash table and *pread()* system call is used to load the record to the memory. *Pread()* allows multiple threads to perform I/O on the same file descriptor in parallel without any locking.

Compression and decompression are implemented using the well-established *zlib* library (DEFLATE algorithm) that is available by default on almost all systems. Each read is compressed/decompressed independently from one another by using an individual *zlib stream* for each read. Thus, multiple reads can be accessed and decompressed in parallel using multiple threads.

### FAST5/SLOW5 conversion, compression and manipulation: slow5tools

*Slow5tools* is implemented on top of the *slow5lib* using C/C++ programming language and follows ISO C++ 2011 standard. Both *slow5lib* and *slow5tools* support Unix systems (Linux and MacOS) or even Windows using the Windows subsystem for Linux. They can be compiled using GNU C/C++ compiler (*gcc/g++*) or LLVM C/C++ compiler (*clang/clang++*) or Intel C/C++ Compiler (*icc/icpc*). We have thoroughly tested both on older systems (e.g., *Ubuntu 14*) as well as modern systems (*Ubuntu 20*). We have tested *slow5lib and slow5tools* on Intel and ARM (both 32 and 64 bit) processors.

### SLOW5 benchmarking experiments

The benchmarking datasets described in **Supplementary Table 1** were generated by sequencing genomic DNA from the human NA12878 reference sample on an ONT PromethION device. Unsheared DNA libraries were prepared using the ONT LSK109 ligation library prep and two flow cells were used to generate ^~^30X genome coverage.

To perform computational benchmarking experiments at realistic workloads, we integrated *slow5lib* to *f5c v0.2* CPU version which is essentially a restructured version of *Nanopolish* that enables us to accurately measure the time for each individual component of a methylation calling job. FAST5 benchmarks were performed using the same version of *f5c* that uses HDF5 1.10.4 built with the *threadsafe* option enabled. POSIX threads are used in *f5c* to perform multithreaded access to FAST5 and SLOW5. To obtain the BAM file for methylation calling, the reads were mapped to the *hg38* reference genome (with no alternate contigs) using *minimap2* version 2.17-r941 (with *-x map-ont -a --secondary=no* options) and sorted using *samtools* v1.9.

Measurement and calculations were performed as follows:

1. The Overall execution time (wall-clock time) and the CPU time (user mode + kernel mode) of the program were measured by running the program through the GNU time utility in Linux.
2. The CPU utilisation percentage is computed as:

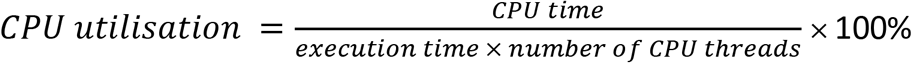

Note that this CPU utilisation percentage is a normalised value based on the number of CPU threads that the program was executed with.
3. Execution time for individual components (I/O operations and data processing) was measured by inserting *gettimeofday()* function calls into appropriate locations in the software source code. To prevent the operating system disk cache affecting the accuracy of I/O results, we cleared the disk cache (pagecache, dentries and inodes) each time before a program execution except on the NCI cluster where this was not permitted. On NCI, disk cache could not be cleaned as we did not have root access, so we implemented a custom program that writes and reads back hundreds of gigabytes of data (several times the size of RAM) to the storage after each experiment so the cache is filled with this mock data. Despite the effect of the hardware disk controller cache (8GB) being negligible due to the large dataset size (>100GB), we still executed a mock program run prior to each experiment.
4. ‘Core-hours’ is calculated as the product of the number of processing threads employed and the number of hours (wall-clock time) spent on the job. This metric is inspired by the metric ‘man-hours’ used in the labour industry and is used in the cloud computing domain to calculate the data processing cost. In an ideally parallel program, this metric remains constant with the number of cores/threads.
5. The disk usage for different files was measured using the *du* command.

## DATA & CODE AVAILABILITY

SLOW5 format and all associated software are free and open source. Datasets used in benchmarking experiments are described in **Supplementary Table 1** and will be hosted in a public repository prior to publication.

SLOW5 format specification documents can be accessed at: https://hasindu2008.github.io/slow5specs *Slow5lib* can be accessed at: https://hasindu2008.github.io/slow5lib/ *Slow5tools* can be accessed at: https://hasindu2008.github.io/slow5tools/

## ACKNOWLEDGEMENTS

We thank our colleagues Derrick Lin, Dmitry Degrave and Warren Kaplan for providing excellent technical support and, most importantly, freedom to use the institute’s HPC system in some quite exotic ways. We thank Leonard Goldstein for critical feedback during manuscript preparation. Resources from the Australian National Computational Infrastructure (NCI) were used during benchmarking experiments. We acknowledge the following funding support: MRFF Investigator Grant MRF1173594 (to I.W.D.) and philanthropic support from The Kinghorn Foundation.

## AUTHOR CONTRIBUTIONS

All authors (H.G., H.Sk., S.P.J., J.M.F., T.G.A., J.M.H., H.Sd., M.A.S., S.P. & I.W.D.) contributed to the conception, design and testing of SLOW5 format. H.G., H.Sk., S.P.J., & J.M.F., implemented SLOW5 format and associated software. J.M.H. generated the ONT sequencing data used in this study. H.G., H.Sk., S.P.J., J.M.F. performed benchmarking experiments. H.G., H.Sk. & I.W.D. prepared the figures. H.G. & I.W.D prepared the manuscript, with support from all authors.

## DISCLAIMERS & COMPETING INTERESTS

I.W.D. manages a fee-for-service nanopore sequencing facility at the Garvan Institute of Medical Research that is a customer of Oxford Nanopore Technologies but has no further financial relationship. H.G., H.Sk., J.M.F., J.M.H. & M.A.S. have received travel and accommodation expenses to speak at Oxford Nanopore Technologies conferences. The authors declare no other competing financial or non-financial interests.

