## Supplementary Materials for "SLOW5: a new file format enables massive acceleration of nanopore sequencing data analysis"

### LIST OF SUPPLEMENTARY MATERIALS

**Supplementary Table 1. Datasets used for benchmarking experiments.**

**Supplementary Table 2. Specifications of all computers used in this study.**

**Figure S1. Inefficient parallel access is a major bottleneck in analysis of FAST5 files.**

**Figure S2. Performance metrics for DNA methylation profiling with FAST5 / SLOW5 files.**

**Figure S3. FAST5 to SLOW5 data conversion performance.**

**Supplementary Note 1. FAST5 format de-mystified.**

**Supplementary Note 2. An inherent limitation in FAST5 files prevents efficient parallel analysis.**

**Supplementary Note 3. SLOW5 format specifications.**

**Supplementary Table 1. Datasets used for benchmarking experiments.**

| Dataset | Platform | Pore | Reads | Total seq. (Gbases) | Size (GB) FASTQ | Size FAST5 | Size SLOW5 | Size BLOW5 | Size compressed-BLOW5 |
|---|---|---|---|---|---|---|---|---|---|
| ~30X human genome (NA12878) | PromethION | R9.4.1 | 9,083,052 | 93.4 | 176 GB | 1.3 TB | 4.0 TB | 2.0 TB | 1.0TB |
| Downsampled human dataset (NA12878) | PromethION | R9.4.1 | 500,000 | 5.1 | 9.5 GB | 71 GB | 212 GB | 106 GB | 53 GB |

**Supplementary Table 2. Specifications of all computers used in this study.**

| System | Type | CPU | CPU cores | RAM (GB) | GPU | File system | Disk System | OS |
|---|---|---|---|---|---|---|---|---|
| HPC-HDD | HPC with HDD RAID | 2 × Intel Xeon Gold 6154 | 36 | 384 | - | ext4 | 12×10TB HDD drives with RAID6 configuration | Ubuntu 18.04.3 LTS |
| HPC-Lustre | NCI CPU node with distributed lustre file system | 2 x core Intel Xeon Platinum 8274 | 48 | 192 | - | lustre | 7200 4TB disks in 120 NetApp disk arrays | CentOS 8.3.2011 |
| HPC-GPU | NCI GPU node with distributed lustre file system | 2 x Intel Xeon Platinum 8268 | 48 | 384 | Nvidia Tesla V100-SXM2-32GB (Volta architecture) | lustre | 7200 4TB disks in 120 NetApp disk arrays | CentOS 8.3.2011 |
| Cloud-FsX | Amazon AWS c5a.16xlarge instance with distributed lustre file system | AMD EPYC 7R32 (virtual machine) | 32 virtual machine | 128 | - | Amazon FSx for Lustre | Amazon FSx Lustre storage (persistent HDD) | Ubuntu 20.04.2 LTS |
| Cloud-EBS | Amazon AWS c5a.16xlarge instance with elastic block storage (EBS) | AMD EPYC 7R32 (virtual machine) | 32 virtual machine | 128 | - | ext4 | 1x500GB elastic block storage (standard magnetic) | Ubuntu 20.04.2 LTS |
| Workstation-SSD | GPU Workstation with SSD | 1xAMD Ryzen Threadripper 3970X | 32 | 128 | NVIDIA 3090 - 24GB (Ampere architecture) | ext4 | 4x500GB SSD drives with RAID0 configuration | Ubuntu 18.04.5 LTS |
| Workstation-NFS | Workstation with network file system (NFS) | 1xAMD Ryzen Threadripper 3970X | 32 | 128 | - | ext4 mounted as NFS | 12x12 TB HDD with RAID 10 configuration (Synology DS3617xs NAS) | Ubuntu 18.04.5 LTS |
| Laptop-HDD | Dell XPS 9570 laptop with a USB 3.0 external HDD attached | Intel core i7-8750H CPU | 6 | 16 | NVIDIA 1050 Ti - 4 GB (Pascal architecture) | ext4 | 1x 500GB HDD (external USB3.0 drive) | Ubuntu 18.04.1 LTS |
| Embedded system | Jetson Xavier AGX with a USB 3.0 external HDD attached | ARM v8 64-bit CPU | 8 | 16 | NVIDIA tegra - 16GB shared with RAM (Volta architecture) | ext4 | 1x 500GB HDD (external USB3.0 drive) | Ubuntu 18.04.3 LTS |

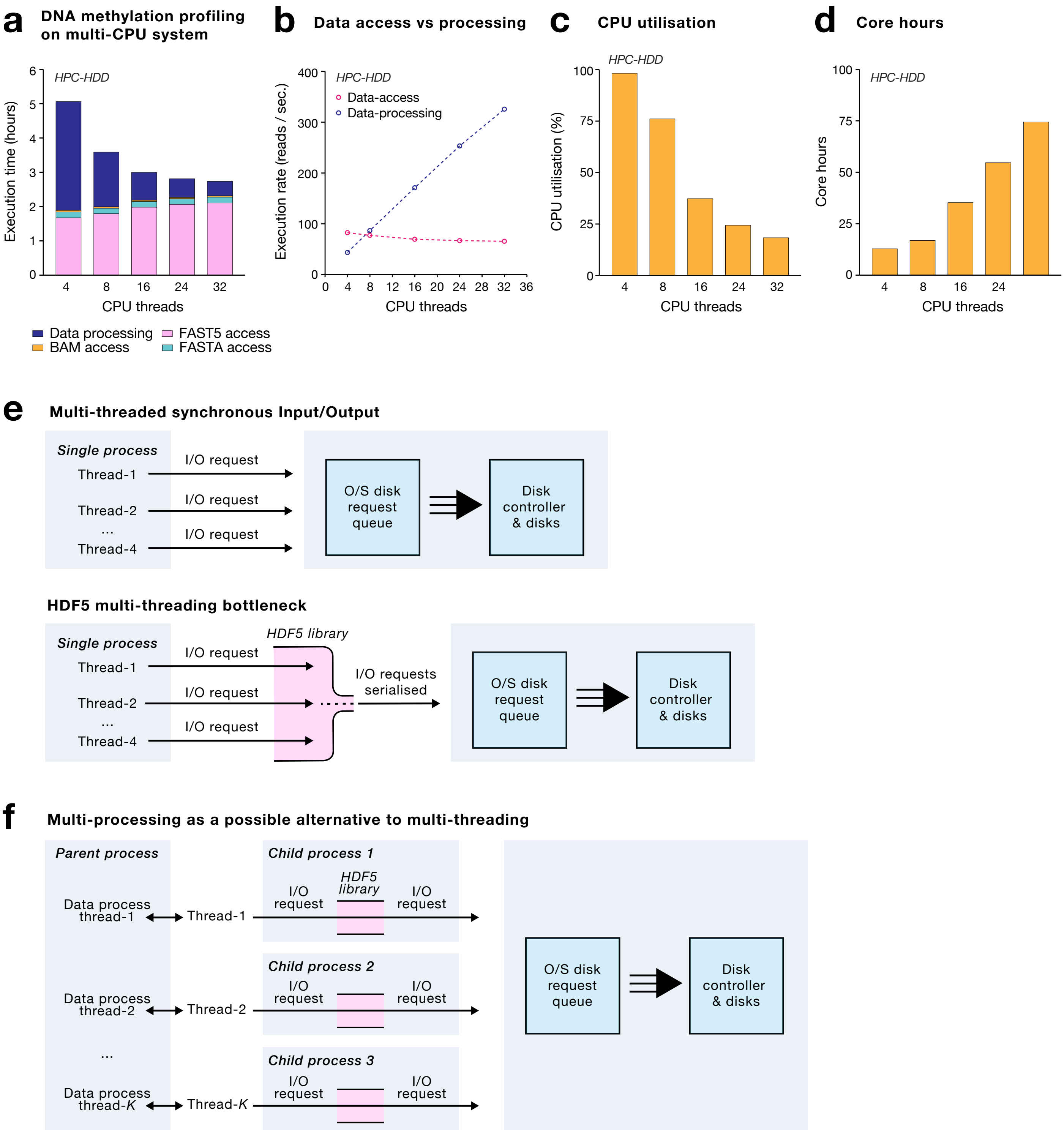


**Figure S1. Inefficient parallel access is a major bottleneck in analysis of FAST5 files.** (**a**) Bar chart shows the time consumed by individual components of a *Nanopolish* DNA methylation calling job with signal data input in FAST5 format: FAST5 data access (pink), FASTA data access (teal), BAM data access (orange) and data processing (navy). To assess the impact of multi-threading, the analysis was run with various numbers of CPU threads on the *HPC-HDD* system (see **Supplementary Table 2**). The analysis was run on a downsampled human genome sequencing dataset of 500 million reads (see **Supplementary Table 1**). (**b**) Dot plots show the rate of file access and processing (reads / second) during the DNA methylation calling job above, as a function of CPU threads used. (**c,d**) Bar charts show the proportional CPU utilisation (**c**) and total core hours (**d**) during the DNA methylation calling job above. The definition of core-hours is provided in the **Methods** section. (**e**) The upper schematic illustrates the architecture of a job with multi-threaded synchronous file access (I/O). The lower schematic illustrates the bottleneck created by the HDF5 library that is required to read FAST5 files. The HDF5 library serialises I/O requests, making multi-threaded analysis highly inefficient and causing the observed decline in CPU utilisation with increasing numbers of CPU threads. (**f**) Schematic illustrates the architecture of a multi-processing approach that was implemented to circumvent this limitation in the HDF5 library. The multi-processing approach is viable, but requires challenging software engineering and is not a generalisable, long-term solution.

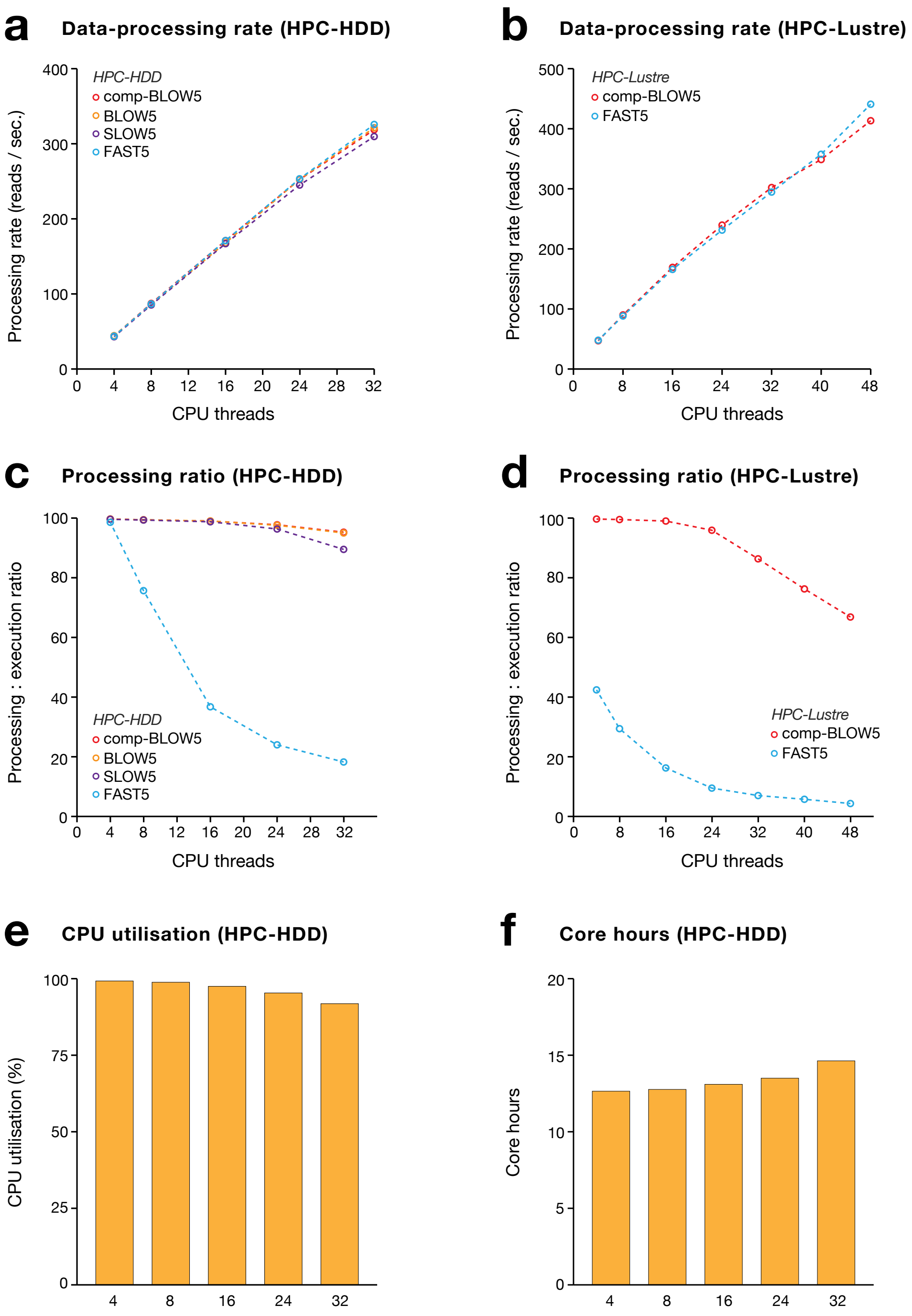


**Figure S2. Performance metrics for DNA methylation profiling with FAST5 / SLOW5 files.** (**a**,**b**) Dot plots show the rate of data processing (reads / second) during *Nanopolish* DNA methylation calling with FAST5 (blue), ASCII SLOW5 (purple), binary BLOW5 (orange) and compressed-BLOW5 (comp-BLOW5; red) files as a function of CPU threads. Analysis was performed on two HPC architectures: *HPC-HDD* (**a**) or *HPC-Lustre* (**b**; see **Supplementary Table 2**). (**c**,**d**) Dot plots show the ratio of data-processing time relative to total execution time for the jobs above. (**e**,**f**) Bar charts show the proportional CPU utilisation (**e**) and total core hours (**f**) during the DNA methylation calling with comp-BLOW5 on the *HPC-HDD* system. The definition of core-hours is provided in the **Methods** section.

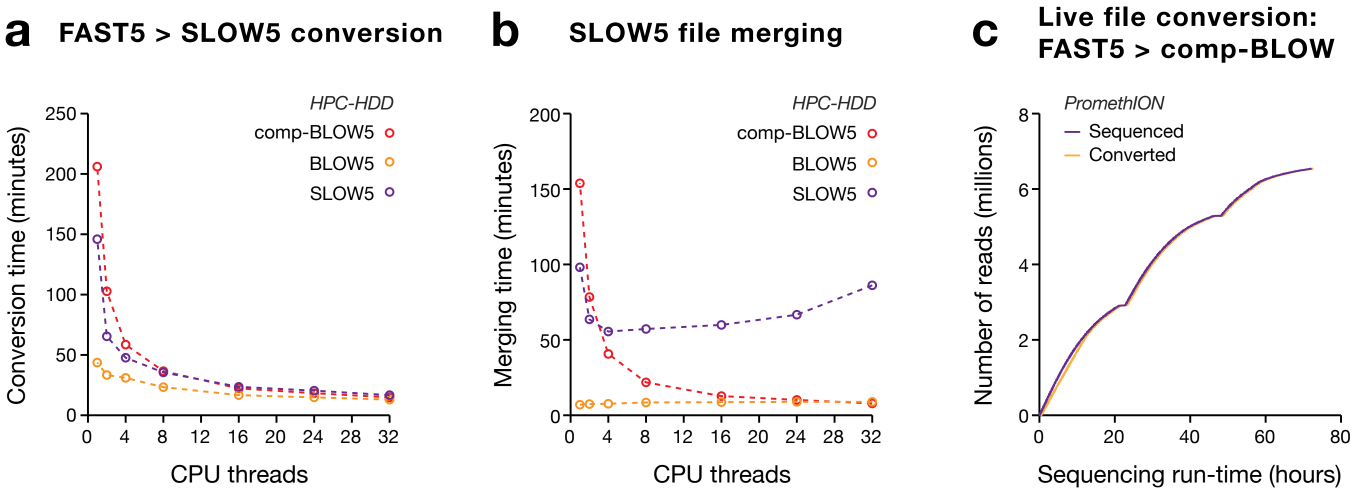


**Figure S3. FAST5 to SLOW5 data conversion performance.** (**a**) Dot plot shows the time take to convert a downsampled human genome sequencing dataset of 500 million reads (see **Supplementary Table 1**) from FAST5 format to ASCII SLOW5 (purple), binary BLOW5 (orange) and compressed-BLOW5 (comp-BLOW5; red) formats as a function of CPU threads used on the *HPC-HDD* system (see **Supplementary Table 2**). (**b**) Dot plot shows the time taken to merge the individual files from (**a**) into a single SLOW5/BLOW5/comp-BLOW5 file. (**c**) Curves show the progress of data generation (purple) and FAST5 to comp-BLOW5 conversion (orange) during a sequencing run on an ONT PromethION device with live conversion enabled. As evident, all reads are converted within minutes of availability and the entire dataset is converted to comp-BLOW5 format at the sequencing run completion.
