## Supplementary Note 1 for "SLOW5: a new file format enables massive acceleration of nanopore sequencing data analysis"

29 June 2021

### Supplementary Note 1. FAST5 format de-mystified

#### PREAMBLE

FAST5 files are Hierarchical Data Format 5 (HDF5) files with a specific schema defined by Oxford Nanopore Technologies (ONT) for storing raw current-signal data generated from ONT devices. To our knowledge, there is no formal, public-facing documentation that fully describes the FAST5 file structure. We have compiled the following material to help researchers to understand FAST5 files. This is based on our own inferences about FAST5 files, and should not be interpreted as a definitive specification document from the developers of FAST5. Please contact us if you are aware of any details we have missed or misinterpreted.

There are two FAST5 types: single-FAST5 and multi-FAST5 (first appearing around September 2018). A multi-FAST5 file contains a batch of reads in a single file whereas a single-FAST5 file contains a single read. Single-FAST5 (one read per file) is no longer used. In this document, FAST5 will always refer to multi-FAST5 unless otherwise stated.

To read FAST5 files we use the HDF5 library and HDF5 tools [1].

#### BASICS

A FAST5 (HDF5) file is like a file system. Just as there are multiple levels of ***directories*** and ***files*** in a file system, a FAST5 (HDF5) file contains ***groups*** and ***datasets***, respectively. The term **HDF5 objects** is an umbrella term for both groups and datasets. HDF5 objects can optionally contain **attributes**, which are key-value pairs. HDF5 related terms are defined below with examples.

##### *Groups*

HDF5 groups (and links[1]) organise HDF5 objects. Every HDF5 file contains a root group that can contain other groups or links to other HDF5 objects[2]. Working with groups and group members (HDF5 objects) is similar in many ways to working with directories and files in UNIX. As with UNIX directories and files, objects in an HDF5 file are often described by giving their full (or absolute) path names.

- */* signifies the root group.
- */foo* signifies a member of the root group called *foo*.
- */foo/zoo* signifies a member of the group *foo*, which in turn is a member of the root group.

##### *Datasets*

HDF5 datasets organise and contain the actual data values [2]. A dataset consists of metadata (datatype, datasize, compression technique, etc) that describes the data, in addition to the data

[1] Links are like directory/file paths in a file system. Links can be absolute, relative or even symbolic.
[2] Other objects can be in the same FAST5 file or in a different FAST5 file (but have not seen this in FAST5 files).

itself. In any read within a FAST5 file, two datasets are found; one inside the *Raw* group (the raw current-signal) and the other inside the *Analyses* group (this is the FASTQ data, if live base-calling was enabled) [1].

##### *Attributes*

Attributes can optionally be associated with HDF5 objects. The attributes have two parts: a name and a value. Attributes are accessed by opening the object that they are attached to; hence, they are not independent objects. Typically an attribute is small in size and contains details about the object that it is attached to.

Attributes look similar to HDF5 datasets in that they have metadata such as data type and dataspace. However, unlike HDF5 datasets, HDF5 attributes do not support partial I/O operations and cannot be compressed or extended [2].

##### *HDF5 dataspace*

The HDF5 *dataspace* must be defined prior to defining an HDF5 dataset or an attribute. The dataspace defines some metadata such as the size and shape of the dataset or attribute raw data, i.e., the number of dimensions and the size of each dimension of the multidimensional array in which the raw data is represented [2].

#### HIERARCHY OF A MULTI-FAST5 FILE

The root group (e.g. /fmh_15… in the snapshot below) contains a group for each read that is named as "read_" followed by the read identifier (e.g. read_001f4… in the snapshot below):

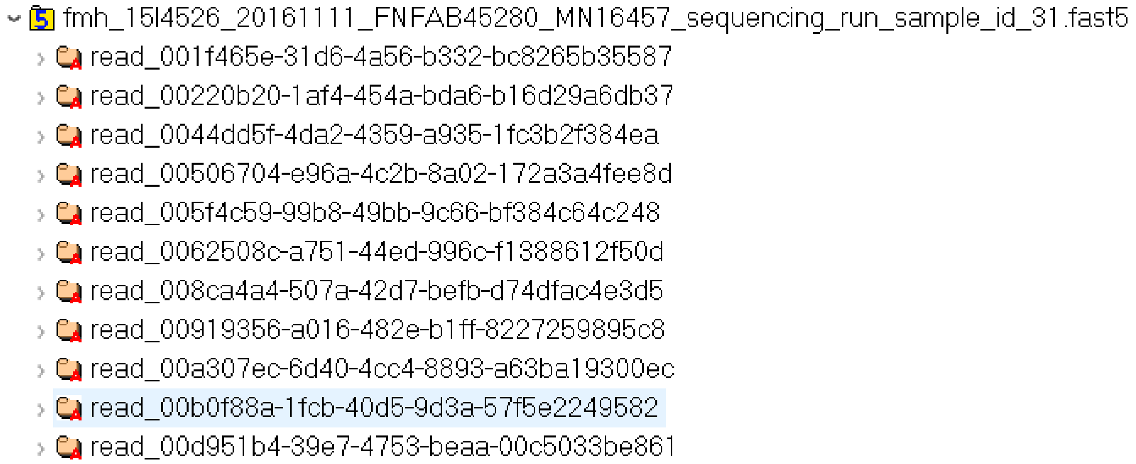

Under each read group, there are the following groups:

- Raw
- channel_id
- context_tags
- tracking_id
- Analyses

Note that the *Analyses* group is only available in base-called FAST5 files. The groups in a FAST5 file that has not been base-called are shown in the snapshot below:

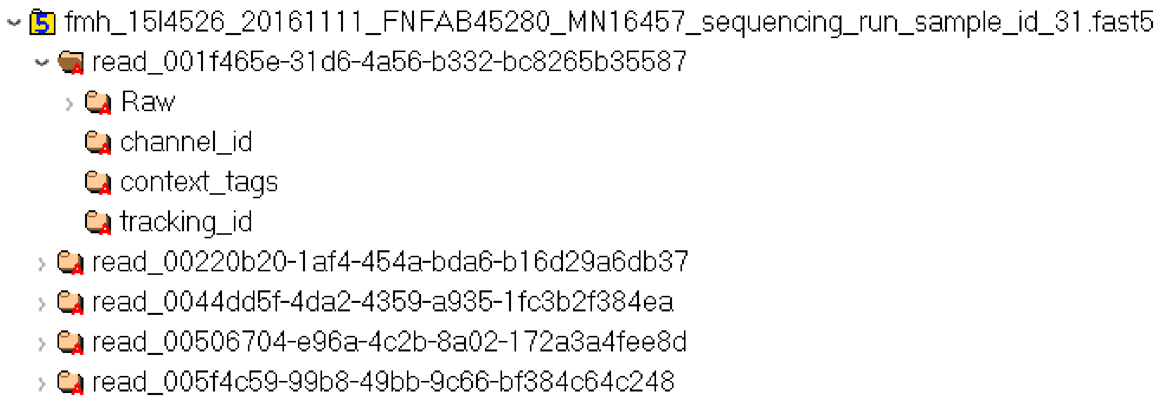

As the name suggests, *Raw* contains the raw signal (raw data acquisition values) and associated metadata. *channel_id* contains (but is not limited to) parameters useful for converting the raw signal into pico-ampere values. *context_tags* and *tracking_id* contain global information that are common to the sequencing run. More information on these groups is provided below.

The *Analyses* group is for storing data resultant from various downstream analyses, such as base-calling. For instance, if the Guppy basecaller is run with the option to output base-called FAST5 files, those output FAST5 files will contain this *Analysis* group. *Analyses* groups can be used by custom software (e.g. Tombo) for storing data from additional downstream analyses.

In the following subsections we provide detailed descriptions of groups mentioned above, except the *Analyses* group. Since we are concerned with the raw signal data, it is not in the scope of this document to discuss that *Analyses* group.

##### ***Root_group***

Root group has two attributes *file_type* (note that *file_type* seems to be only available in multi-fast5 version 2.2) and *file_version*. (Note that ONT's version control scheme is somewhat inconsistent, so we make no assumptions about file-structure based on FAST5 version numbers).

1. **file_type**

   *Example value: multi-read*

   *Data type: String*

2. **file_version**

   *Example value: 2.2*

   *Data type: String*

##### ***Read***

A read group has two attributes *run_id* and *pore_type* (note that *pore_type* is not available in multi-fast5 v2.0).

1. **run_id**

   The value of this attribute is constant across all the read groups.
   *Example value: fe697f519ab04ba540bc4fe93f7cbd86669f38ca*
   *Data type: String*
2. **pore_type**

   In existing FAST5 versions, this value is empty. This attribute may be used in the future to distinguish different pore types within a single flow cell.
   *Example value: <not set>*
   *Data type: String*

#### *Raw*

*Raw* group contains one dataset and seven attributes. The dataset is the raw signal which is a series of 16-bit integers (HDF5 Datatype = *H5T_STD_I16LE*). These are the integer values directly coming from the data acquisition process (analog to digital converter). This raw signal can be converted into pico Ampere (pA) values using attributes available in the *channel_id* group (explained later).

The seven attributes from the *Raw* group are listed below with a description of each attribute, example values and the data type. Understanding these descriptions require a brief understanding of an ONT flow cell. A flow cell has multiple channels allowing multiple DNA/RNA strands to be sequenced in parallel. For instance, a MinION flow cell has 512 channels and thus can sequence 512 strands in parallel. Each channel contains one or more wells[3]. For instance, a MinION flow cell has 4 wells per channel. The wells within a channel are connected to a multiplexer (MUX), a switch that controls which of the four wells in the channel is controlled and read out by the circuits. Please refer to reference [3] or [4] for more information about channels and multiplexers.

Note that some of the information below is extracted from ONT document [1].

1. **read_number**

   This number is unique within the channel, but only really meaningful in respect to the bulk FAST5[4] file read table
   *Example value: 17981*
   *Data type: 32-bit unsigned integer*
2. **read_id**

   A unique identifier for the read. This is a Universally unique identifier (UUID), and should be unique for any read from any device
   *Example value:* 00592138-f120-4ab5-9916-c5567adb8e29
   *Data type: String*
3. **start_time**

[3] Each well should ideally contain one pore
[4] ONT provides an optional bulk FAST5 file format to capture the entire data stream from every channel on the sequencing device. This file includes raw signal and metadata for every channel including MinKNOW classifications

The start time of the read. The unit for *start_time* is 'number of sampling events', so *start_time* has to be divided by sampling rate (Read_xxxx/channel_id/sampling_rate) to get the start time in seconds (i.e. the time since the run was started).

*Example value:* 335845487

*Data type: 64-bit unsigned integer*

4. **duration**

The duration of the read. The unit for *duration* is also 'number of sampling events'.

*Example value:* 335845487

*Data type: 32-bit unsigned integer*

5. **start_mux**

The MUX setting[5] for the channel when the read began. Due to timing issues this can sometimes reflect what the MUX was just *before* the read began; this will only matter for reads that start immediately after a MUX change.

*Example value: 4*

*Data type: 8-bit unsigned integer*

6. **median_before**

The measure of the median current level for the data preceding the read. In most cases this can be used as an estimate of the open pore level[6].

*Example value:* 238.78225708007812

*Data type: 64-bit floating-point*

7. **end_reason**

This is a new attribute in FAST5 v2.2 that is currently not stable (i.e. it is present in some v2.2 files but not others).

*Example value:* unblock_mux_change

*Data type: 8-bit enum*

```
ATTRIBUTE "end_reason" {
    DATATYPE  H5T_ENUM {
    H5T_STD_U8LE;
    "unknown"       0;
    "partial"       1;
    "mux_change"            2;
    "unblock_mux_change" 3;
    "signal_positive"  4;
    "signal_negative"  5;
    }
```

#### *Channel_id*

This group has attributes that are relevant to the channel that sequenced a given read. The channel_id group has the following attributes.

1. **channel_number**

*Example value:* 504

[5] out of the wells in the channel, which well the mux is set to sequence

[6] open-pore state is when there is no strand inside the pore

*Data type: String*

2. **digitisation**

   The digitisation is likely to be the number of quantisation levels in the Analog to Digital Converter (ADC). That is, if the ADC is 12 bit, digitisation is 4096 (2^12).

   *Example value:* 8192.0

   *Data type: 64-bit floating-point*

3. **offset**

   Likely to be the ADC offset error. This value is added when converting the signal to pico ampere.

   *Example value:* 10.0

   *Data type: 64-bit floating-point*

4. **range**

   Likely to be the full scale measurement range in pico amperes.

   *Example value:* 1441.389892578125

   *Data type: 64-bit floating-point*

5. **sampling_rate**

   Sampling frequency of the ADC, i.e., the number of data points collected per second.

   *Example value:* 4000

   *Data type: 64-bit floating-point*

Of these attributes, *digitisation*, *offset* and *range* can be used to transform the raw signal in the *Raw* group, to pico-ampere current values as follows:

```
signal_in_pico_ampere = (raw_signal_value + offset) * range / digitisation
```

#### Context_tags

The *context_tags* group has global attributes that describe the sequencing run. The attributes under the *context_tags* group are listed below with short descriptions. (Note that these descriptions are mostly inferred from our own experiences and may not be totally correct). The data type of the value of all the attributes listed under *context_tags* group is String.

1. **barcoding_enabled**

   Indicates if barcode demultiplexing is enabled during live basecalling

   *Example value:* 0

2. **experiment_duration_set**

   Indicates the duration of the experiment selected when starting the sequencing run (likely to be in minutes)

   *Example value:* 4320

3. **experiment_type**

   Indicates the type of the experiment, for instance, *genomic_dna* or *rna*.

   *Example value:* genomic_dna

4. **local_basecalling**

   Indicates if live base calling is enabled or not.

   *Example value:* 1

5. **package**

   We are note sure about this attribute, but it seems to relate to Bream [https://github.com/nanoporetech/minknow_lims_interface]

   *Example value:* bream4
6. **Package_version**

   *Example value:* 6.0.7
7. **sample_frequency**

   Seems to be the same as the *sampling_frequency* in the *channel_id* group

   *Example value:* 4000
8. **sequencing_kit**

   The sequencing kit used, for instance, *sqk-lsk109* or *sqk-rna002*. [https://store.nanoporetech.com/sample-prep.html]

   *Example value:* sqk-lsk109

There can be additional attributes such as *basecall_config_filename*, depending whether live basecalling was turned on/off when the sequencing run was started.

#### *Tracking_id*

The *tracking_id* group has global attributes relevant to the sequencing run and the sequencing device. Much of this is mysterious to us, but likely used internally by ONT. The data type of the value of all the attributes listed under *tracking_id* group is String.

1. **asic_id**

   Possibly the Application Specific Integrated Circuit identifier (ASIC) of the flowcel (unique number of the chip), for nanopore tracking. Nanopore presumably uses this to see good/bad batches of chips.

   *Example value:* 213553007
2. **asic_id_eeprom**

   Possibly the identifier of the ASIC's electrically erasable programmable read-only memory (EEPROM) of the flow cell.

   *Example value:* 5309577
3. **asic_temp**

   The temperature in degrees celsius of the ASIC chip presumably at the start of the sequencing run.

   *Example value:* 28.867193
4. **asic_version**

   The version of ASIC being used

   *Example value:* IA02D
5. **auto_update**

   Possibly whether auto update in Minknow is enabled or not.

   *Example value:* 0
6. **auto_update_source**

   The link to the Minknow update source.

   *Example value:* https://mirror.oxfordnanoportal.com/software/MinKNOW/

7. **bream_is_standard**
   Bream is one of the software for controlling sequencing. So something to do with that.
   *Example value:* 0
8. **configuration_version**
   Probably the MinKNOW core version .
   *Example value:* 4.0.13
9. **device_id**
   The serial ID of the MinION or device position for GridION/PromethION.
   Device position on GridION/PromethION refers to the ID of the bay (slot where the flowcell is put) on the device.
   *Example value:* X2
10. **device_type**
    The device type, that is whether MinION, PromethION or GridION
    *Example value:* gridion
11. **distribution_status**
    Stable vs dev/alpha/beta status.
    *Example value:* stable
12. **distribution_version**
    MinKNOW version maybe.
    *Example value:* 20.06.9
13. **exp_script_name**
    These are the scripts used based on what kits are selected in MinKNOW for sequencing.
    *Example value: sequencing/sequencing_MIN106_DNA:FLO-MIN106:SQK-LSK109*
14. **exp_script_purpose**
    Likely to be if a real sequencing run or a simulation playback.
    *Example value:* sequencing_run
15. **exp_start_time**
    Start time of sequencing run.
    *Example value:* 2020-09-08T01:23:21Z
16. **flow_cell_id**
    Unique ID for the flowcell, used by ONT to track flowcell metrics and warranty.
    *Example value: FAN43349*
17. **flow_cell_product_code**
    The type of flowcell, these will be different based on R9.4.1, R10.3, R9.5, PromethION, etc.
    *Example value: FLO-MIN106*
18. **guppy_version**
    Guppy version being used by MinKNOW.
    *Example value: 4.0.11+f1071ce*
19. **heatsink_temp**
    The heat sink that is on the sequencer that is used to control the ASIC temp in degrees celsius most probably at the start of the sequencing run.
    *Example value: 33.996094*

20. **hostname**
    The hostname of the computer doing the sequencing run.
    *Example value: GXB02243*
21. **installation_type**
    This is the MinKNOW install type. We believe that for GridION and MinION it is nc, and something else for PromethION.
    *Example value: nc*
22. **local_firmware_file**
    *Example value: 1*
23. **operating_system**
    The operating system and the version of the computer performing the sequencing run.
    *Example value: ubuntu 16.04*
24. **protocol_group_id**
    This is the run name given by the user for the experiment group. We could have multiple run-ids if we run multiple flowcells under the same experiment "group" name.
    *Example value: GLFN180082*
25. **protocol_run_id**
    I think this is a unique UUID for the experiment GROUP (just in case the name given by the user is not unique). This is same for each run of the same experimental group.
    *Example value: f2c69573-5fef-43b8-8d81-9cb20634aa7c*
26. **protocols_version**
    Not sure exactly what this is, but probably a way MinKNOW tracks it's various protocols for barcoding, kits, etc.
    *Example value: 6.0.7*
27. **run_id**
    The unique run ID which will be different for each run, even in the same experiment group.
    *Example value: 07770780274b0e3703f00d969291b1a37a5a6be1*
28. **sample_id**
    Sample ID is the name given by the user for the sample.
    *Example value: NA12878*
29. **usb_config**
    Various information about the connection between the flowcell and the computer.
    *Example value: GridX5_fx3_1.1.3_ONT#MinION_fpga_1.1.1#bulk#Auto*
30. **version**
    *Example value: 4.0.3*

#### FAST5 VERSIONS & THEIR ATTRIBUTES

The following table shows the availability (and unavailability) of attributes in un-basecalled multi-FAST5 files for a file with versions 2.0 and another file with version 2.2. We could not get hold of any files from version 2.1, but please contact us if you are aware of such a version.

29 June 2021

Green cells = attribute available.

| Group | Attribute name | V2.0 | V2.2 |
|---|---|---|---|
| / | file_type | | green |
| | file_version | green | green |
| /read | run_id | green | green |
| | pore_type | | green |
| /read/Raw | start_time | green | green |
| | duration | green | green |
| | read_number | green | green |
| | start_mux | green | green |
| | read_id | green | green |
| | median_before | green | green |
| | end_reason | | green |
| /read/channel_id | digitisation | green | green |
| | offset | green | green |
| | range | green | green |
| | sampling_rate | green | green |
| | channel_number | green | green |
| /read/context_tags | barcoding_enabled | | green |
| | experiment_duration_set | | green |
| | experiment_type | | green |
| | local_basecalling | | green |
| | package | | green |
| | package_version | | green |
| | sample_frequency | green | green |
| | sequencing_kit | | green |
| | experiment_kit | green | |
| | filename | green | |
| | user_filename_input | green | |
| /read/tracking_id | asic_id | green | green |
| | asic_id_eeprom | green | green |
| | asic_temp | green | green |
| | asic_version | | green |
| | auto_update | green | green |
| | auto_update_source | green | green |
| | bream_core_version | green | |
| | bream_is_standard | green | green |
| | bream_ont_version | green | |
| | bream_prod_version | green | |
| | bream_rnd_version | green | |

| | |
|---|---|
| | configuration_version |
| | device_id |
| | device_type |
| | distribution_status |
| | distribution_version |
| | exp_script_name |
| | exp_script_purpose |
| | exp_start_time |
| | flow_cell_id |
| | flow_cell_product_code |
| | guppy_version |
| | heatsink_temp |
| | hostname |
| | installation_type |
| | local_firmware_file |
| | operating_system |
| | protocol_group_id |
| | protocol_run_id |
| | protocols_version |
| | run_id |
| | sample_id |
| | usb_config |
| | version |

#### CONSTANT & VARIABLE ATTRIBUTES

Many FAST5 attributes are identical amongst all the reads within a single sequencing run (within multi-FAST5 files as well as amongst different multi-FAST5 files). For example, all reads from a given experiment will have the same *run_id*. Some attributes are variable between different reads, even within a single multi-FAST5. For example, each read has a different *read_id*. All the attributes in *contex_tags* and *tracking_id* are constant across all reads in a single sequencing run, whereas most of the attributes in *Raw* and *channel_id* are variable between reads (with a few exceptions).

The variable attributes amongst all groups (except the *Analyses* group) are:

- duration
- end_reason (not in version 0.6 but in 2.2)
- median_before
- read_id
- read_number
- start_mux
- start_time
- channel_number

- offset

Note that the dataset “Signal” obviously has variable data. All the other attributes are contant.

### ADVANCED INFORMATION

#### *Symbolic links*

For a given FAST5 file, the values of the attributes belonging to the two groups, *context_tags* and *tracking_id* are the same for all the reads in that FAST5 file. Hence only the first read_xxxx group has the actual attributes. The rest of the read_xxxx groups maintain symbolic links [2] to the first read_xxxx group. One can observe the linking structure of a FAST5 file using a utility program called *h5dump* developed by the HDF5 group.

The following is an example output for a FAST5 file where read_000200a4-0347-4a49-b800-37ad7b4287c9 is the first read group. As listed below the rest of the read groups’ *context_tags* and *tracking _id* attributes maintain links (symbolic links) to the *context_tags* and *tracking_id* attributes of the first read group respectively. This observation was valid for all the FAST5 files we have examined.

```
GROUP "context_tags" {
        HARDLINK "/read_000200a4-0347-4a49-b800-37ad7b4287c9/context_tags"
        }
 GROUP "tracking_id" {
        HARDLINK "/read_000200a4-0347-4a49-b800-37ad7b4287c9/tracking_id"
        }
```

#### *ONT h5 validator*

Ont_h5_validator is a tool developed by Oxford Nanopore Technologies to check if a given FAST5 file complies with the FAST5 schema. This tool only considers a subset of the complete FAST5 schema to validate a file.

#### *Single-FAST5 format fields*

The groups, attributes and some example values for a single-FAST5 file are given for the sake of completeness, despite not being used today.

| |
|---|
| **PreviousReadInfo**<br>previous_read_id = cf435984-627d-450d-a81d-2a55c6060c80<br>previous_read_number = 80 |
| **Read**<br>duration = 30695<br>median_before = 206.2032470703125<br>read_id = b3d473e9-34f0-4ad6-a030-61ba6ab458bc<br>read_number = 99<br>start_mux = 4 |

start_time = 318648

**channel_id**
channel_number = 707
digitisation = 2048.0
offset = -196.0
range = 748.5801660113588
sampling_rate = 4000.0

**context_tags**
experiment_duration_set = 3840
experiment_type = genomic_dna
fast5_output_fastq_in_hdf = 1
fast5_raw = 1
fast5_reads_per_folder = 4000
fastq_enabled = 1
fastq_reads_per_file = 4000
filename = pct0028_20181029_0004a30b00232bec_1_e11_h11_sequencing_run_lxbab132606_84140
flowcell_type = flo-pro002
kit_classification = none
local_basecalling = 1
local_bc_comp_model =
local_bc_temp_model = template_r9.4_450bps_5mer_raw.jsn
sample_frequency = 4000
sequencing_kit = sqk-lsk109
user_filename_input = lxbab132606

**tracking_id**
asic_id = 0004A30B00232BEC
asic_id_eeprom = 0004A30B00232BEC
asic_temp = 36.990513
asic_version = Unknown
auto_update = 0
auto_update_source = https://mirror.oxfordnanoportal.com/software/MinKNOW/
bream_is_standard = 0
device_id = 1-E11-H11
device_type = promethion
exp_script_name = 59dfa94107ee2b6c0f4be0822482e7da35b4116a-da65898430ab8c4bfe54ba7064f0301390b76211
exp_script_purpose = sequencing_run
exp_start_time = 2018-10-29T01:40:23Z
flow_cell_id = PAD11989
heatsink_temp = 41.996017
hostname = PCT0028
hublett_board_id = 013220e36be4c748
hublett_firmware_version = 2.0.5
installation_type = nc
ip_address =
local_firmware_file = 1
mac_address =
operating_system = ubuntu 16.04
protocol_run_id = e3b445eb-5626-48ef-acc2-b28bcc611009
protocols_version = 0.0.0.0
run_id = 855cdb4b269484b72699b681e539e090c4a50bbb
sample_id = LXBAB132606
satellite_board_id = 0000000000000000
satellite_firmware_version = 2.0.4
usb_config = firm_1.2.3_ware#rbt_4.5.6_rbt#ctrl#USB3
version = 1.14.2
