## Supplementary Note 2 for "SLOW5: a new file format enables massive acceleration of nanopore sequencing data analysis"

### Supplementary Note 2. An inherent limitation in FAST5 files prevents efficient parallel analysis.

#### PREAMBLE

High Performance Computing (HPC) systems offer significant computational power through many-core CPUs that can be utilised in parallel. Moreover, HPC systems have Redundant Arrays of Independent Disks (RAID) storage composed of many disks for higher I/O throughput. Multi-threaded analysis on HPC systems is now standard practice in genomics, enabling efficient analysis of large DNA sequencing datasets.

In our experience, the analysis of nanopore signal data (FAST5 files) is generally slow, even on powerful HPC systems. To understand why, we undertook a detailed investigation of a typical signal-level ONT analysis on a typical HPC system. We selected DNA methylation (5mC) profiling with the popular software *Nanopolish* [1] as our example use-case to assess computational performance and identify potential bottlenecks.

#### APPROACH

We executed the *call-methylation* tool within the *Nanopolish* toolkit on a downsampled ONT human genome sequencing dataset of 500 million reads (see **Supplementary Table 1**). We used a restructured version of the *Nanopolish* software that allowed us to record the wall-clock time spent on I/O operations and data processing. This enables the time consumed by individual components of the analysis (FAST5 files access, FASTA file access, BAM file access & data processing) to be monitored separately. The experiment was run on an HPC system 12 × 10TB HDD drives with RAID6 configuration (**Supplementary Table 2**), using either 4, 8, 16, 24 or 32 CPU threads.

#### AN I/O BOTTLENECK LIMITS PERFORMANCE

We observed a relatively modest improvement in the overall execution time with increasing numbers of CPU threads and almost no improvement beyond 16 threads (**Fig. S1a**). While we observed a linear increase in the rate of data processing with additional CPU threads, there was no increase in the rate of FAST5 data access and, hence, FAST5 data access came to represent an increasingly large fraction of the total execution time (**Fig. S1a,b**). We observed a steep decline in CPU utilisation with increasing numbers of threads, dropping to just 18% utilisation at 32 threads (**Fig. S1c**). Likewise, we found that core-hours (which should be constant in an ideal scenario; see definition under **Methods**) increased with additional threads (**Fig. S1d**).

These results clearly demonstrate that additional CPU threads are not efficiently utilised by *Nanopolish* to improve the overall execution time. There can be two possible explanations for the inability of a tool to efficiently utilise parallel resources: (i) a bottleneck in data processing; and (ii) a bottleneck in

Input/Output (I/O). Our observations strongly suggest that an I/O bottleneck is the primary reason for the under-utilisation during *Nanopolish* analysis.

#### UNDERSTANDING THE BOTTLENECK

We employed performance monitoring and profiling tools during the above experiment, in order to elucidate the causes of inefficient resource-utilisation and performance.

*Hypothesis-1: The performance of the software tool is bounded by file I/O.*

We observed through the *htop* utility in Linux that the majority of *Nanopolish* threads were in the 'D' state during the experiment. The 'D' state is defined as the 'state of the process for disk sleep (uninterruptible)'. This suggests that the software is bounded by file I/O.

*Hypothesis-2: The file I/O bottleneck is caused by the HDF5 library and not by the limitation of physical disks.*

We observed disk usage statistics using the *iostat* utility and found that the disk system was not fully utilised during the experiment (i.e., the observed number of disk reads per second was around 100 Input/output operations per second (IOPS)), while the particular disk system could handle more than 1000 IOPS). This implies that the I/O bottleneck is not due to the limitation of physical disks to serve data fast enough to saturate the processor.

To investigate further, we profiled *Nanopolish* with *Intel Vtune* under concurrency profiling. It reveals that the majority of the 'wait time' is due to a conditional variable (synchronisation primitive) in the underlying HDF5 library that is used to access FAST5 files. Closer inspection of the HDF5 library revealed that the thread-safe version of the HDF5 library serialises the calls for disk read requests. Thus, we reasoned that CPU under-utilisation is caused by the disk requests being serialised by the HDF5 library, consequently causing a bottleneck that limits the utility of a multi-disk RAID system.

#### A HDF5 LIBRARY LIMITATION PREVENTS PARALLEL ACCESS

The HDF5 library required to read and write FAST5 files uses synchronous I/O calls and even the latest HDF5 implementation (HDF5-1.10) does not support asynchronous I/O[1]. This, by itself, is not an issue as multiple synchronous I/O operations can be performed in parallel using multiple I/O threads to exploit the high throughput of RAID systems (**Fig. S1e**; upper).

However, the HDF group (that maintains the HDF5 library) mentions that the thread-safe version of the HDF5 library is not thread efficient and that it effectively serialises the calls for disk read requests [2]. The global lock in the thread safe version of the HDF5 library creates limitations, as explained in the following extract from the HDF5 documentation:

*"Users are often surprised to learn that (1) concurrent access to different datasets in a single HDF5 file and (2) concurrent access to different HDF5 files both require a thread-safe version of the HDF5 library. Although each*

[1] In synchronous I/O calls, the OS, upon receiving the call, puts the user-space thread to sleep and the thread can no longer submit I/O requests until the disk reading is completed and woken by the OS. Conversely, asynchronous I/O system calls return immediately without the thread being put to sleep and the thread can continue to submit another asynchronous request.

*thread in these examples is accessing different data, the HDF5 library modifies global data structures that are independent of a particular HDF5 dataset or HDF5 file. HDF5 relies on a semaphore around the library API calls in the thread-safe version of the library to protect the data structure from corruption by simultaneous manipulation from different threads. Examples of HDF5 library global data structures that must be protected are the freespace manager and open file lists."*

Thus, in spite of having multiple I/O threads, I/O requests for HDF5 files have to go through the HDF5 library (**Fig. S1e**; lower). In a scenario where multiple I/O threads are requesting I/O from the HDF5 library in parallel, the lock inside the HDF5 libraries serialises the parallel requests, effectively issuing only one request at a time to the operating system disk request queue.

### A MORE DETAILED EXPLANATION

**Fig. S1e** (upper) illustrates how multiple I/O threads can be used to perform parallel disk accesses using synchronous I/O. Suppose the disk system has *K* disks, up to *K* requests may be served simultaneously depending on the RAID level; i.e., *K* simultaneous parallel reads are possible on a RAID 0 system with *K* disks. Let $t$ be the average disk request service time (from the time of the system call to when the thread is woken up).

For a program that launches *K* I/O threads and if the disk controller can serve *K* requests in parallel, the total time for *n* disk reads is $T' = t \times \frac{n}{K}$.

However, in spite of having multiple I/O threads, I/O requests for HDF5 files have to go through the HDF5 library. **Fig. S1e** (lower) illustrates this, where *K* I/O threads are requesting I/O from the HDF5 library in parallel. However, the lock inside the HDF5 serialises the parallel requests, effectively issuing only one request at a time to the operating system disk request queue. The operating system will put the thread to sleep and this is equivalent to a single I/O thread. Thus, the total time spent on disk accesses *T* will be $T = t \times n$, and essentially, the high throughput capability of multiple disks in a RAID configuration is under-utilised.

### POSSIBLE WORKAROUNDS

We explored several possible approaches to circumvent the FAST5 bottleneck and these are articulated below. However, due to limitations in each of these approaches we decided instead to create an alternative file format that is not dependent on the HDF5 library.

##### *Fixing HDF5 Library*

One possible solution to the limitation articulated above is to re-engineer the HDF5 library to be thread efficient. However, the HDF5 library is a complicated library with a large code base of >300,000 lines of C code and such a fix would need to be carried out by the HDF Group. The HDF5 Group mentions that the future plan to implement efficient multi-threaded access is currently hindered by inadequate resources [2]. Therefore, such a fix is unlikely to happen in the near future. There is no other alternate library to read HDF5 files, including FAST5 files [3, 4].

##### *Using a process pool*

Multiple threads in a single process share the same address space and thus the lock in the HDF5 library affects multiple threads. Multiple threads are typically used to run sub-tasks in parallel while conveniently sharing data amongst the threads. In contrast, multiple processes have their own independent address spaces and are typically used to run isolated tasks in parallel. The presence of independent address spaces in multiple processes can be exploited to circumvent the lock in the HDF5 library.

A multi-process based solution is elaborated in **Fig. S1f**. Multi-threads in the single parent-process are used for data processing and multiple child-processes for I/O. The parent-process performs data processing using multiple threads in parallel. Each child-process has its own instance of the HDF5 library, as a consequence of independent address spaces. Moreover, each child-process has only a single thread that requests I/O. Thus, a single instance of the HDF5 library gets only one request at a time. In effect, there are multiple instances of the HDF5 library that can submit multiple I/O requests in parallel to the operating system (as opposed to the situation in multi-threaded HDF5 case), thus benefiting from the high throughput offered by RAID configurations.

Formally, if there are *K* processes and if the disk controller can serve *K* requests in parallel, the total time spent on I/O operations will be $T' = t \times \frac{n}{K}$.

Multiple processes are spawned at the beginning of the program using the *fork* system call. These forked child-processes form a pool of processes that exist until the lifetime of the parent-process, solely performing I/O of FAST5 files. The data processing can be performed by multiple threads spawned by the parent-process as usual. The parent-process, when it requires to load signal data of *N* reads (FAST5 accesses), first splits the list of reads into *K* parts where *K* is the number of child-processes. Then, each part is assigned to a child-process, which performs the assigned FAST5 accesses. When the data is loaded, the child-processes send this data to the parent-process.

*Note 1:* A fork-join model for multi-processes (as could be done for multi-threading) is unsuitable to be used instead of the process pool model presented above. Firstly, creating a process can be very expensive and could easily become the biggest bottleneck than the file reading itself. Secondly, forking in the middle of a program could double the memory usage and is usually problematic, and should be avoided where possible.

*Note 2:* It is important to note that *processes* in an operating system are meant for isolation whereas *threads* are for sharing data. Inter-process communication requires system calls, while inter-thread communication involves sharing the same memory space. Further, spawning multiple processes is expensive and is not lightweight (unlike threads). Thus, using processes as a replacement to threads makes the code relatively complicated.

In summary, a process pool approach, whilst technically viable, requires complicated re-engineering for every piece of software and is not a geralisable, long-term solution.

***Naive Approaches of Multi-processing***

Instead of using a process pool solely for FAST5 I/O and multi-threads for parallel data processing, developers may use multi-processes for both the I/O operations and parallel data processing. This would be easier than implementing a pool of processes, however, this is only suitable for perfectly parallel cases. This is the method we use in *slow5tools* for fast conversion from FAST5 to SLOW5. If the application needs to share data among multiple processing units, processes are unsuitable due to the complexity that arises when performing inter-process communication.

Alternatively, the developer may let users manually split data and launch multiple processes. Unfortunately, this method exerts additional burden on the user, i.e., custom scripts must be written for data splitting, launching data processing and concatenating the result. Moreover, this is only suitable for perfectly parallel applications where data can be easily split. Also, an expensive HPC system with dozens of cores is superfluous as the user could use a cluster of low cost networked computers [5].
