## Supplementary Note 3 for "SLOW5: a new file format enables massive acceleration of nanopore sequencing data analysis"

### Supplementary Note 3. SLOW5 Specifications (version 0.1.0)

#### PREAMBLE

SLOW5 is a new file format encoding signal data from Oxford Nanopore Technologies (ONT) devices. SLOW5 was developed to overcome inherent limitations in the existing FAST5 (HDF5) data format that prevent efficient parallel analysis and cause many headaches for developers.

SLOW5 refers to two file formats, namely SLOW5 ASCII and SLOW5 binary (called BLOW5). The extension for SLOW5 ASCII is *.slow5* and for BLOW5 it is *.blow5*. For efficient data accessing and to minimise disk space, users are expected to use BLOW5. SLOW5 ASCII is the human readable format of *BLOW5* and should only be used to view the content. Random access to either SLOW5 ASCII or BLOW5 is supported using a binary index file; this is a separate file in the same directory as the SLOW5 ASCII or BLOW5 file. For SLOW5 ASCII, the index takes the extension *.slow5.idx* and for BLOW5 the index takes *.blow5.idx*.

A SLOW5 file contains a header followed by the ONT sequencing data. In ONT datasets, the *run_id* is a unique identifier that distinguishes a sequencing run. We will refer to a sequencing run and its data as a *read group*. A SLOW5 file can store multiple read groups in a single file, allowing data from multiple sequencing runs to be stored in a single SLOW5 file whilst retaining their individual metadata.

Full specifications for current and previous versions of SLOW5 are available at: https://hasindu2008.github.io/slow5specs/

#### SLOW5 ASCII

A SLOW5 ASCII file is a plain text file that uses the American Standard Code for Information Interchange (ASCII) encoding (locale: C/POSIX, code set: US-ASCII). The file extension is *.slow5.*

An example format of a SLOW5 ASCII file with a single read group is provided in **Table 1.** An example format with multiple read groups - i.e., multiple sequencing runs - is provided in **Table 2**. The column/row borders and cell colours are added only to increase the readability. The actual format uses tabs ('\t') and newlines ('\n') as delimiters (IMPORTANT: '\r' or "\r\n" are not allowed). The first set of lines is the SLOW5 header. The header lines start with '#' or '@'. The remainder of the file encodes ONT data in one read per line.

**Table 1: Example of a SLOW5 ASCII file with a single read group.**

Blue = global header. Yellow = data header. White = data records.

| | | | | | | | | |
|---|---|---|---|---|---|---|---|---|
| #slow5_version | 1.0.0 | | | | | | | |
| #num_read_groups | 1 | | | | | | | |
| @asic_id | 0004A30B00232BEC | | | | | | | |
| @exp_start_time | 2020-01-01T00:00:00Z | | | | | | | |
| @flow_cell_id | FAH00000 | | | | | | | |
| @run_id | 855cdb | | | | | | | |
| …. | …. | | | | | | | |
| #char* | uint32_t | double | double | double | double | uint64_t | int16_t* | ... |
| #read_id | read_group | digitisation | offset | range | sampling_rate | len_raw_signal | raw_signal | ... |
| read0 | 0 | 8192 | 6 | 1467.6 | 4000 | 123456 | 498,492,... | ... |
| read1 | 0 | 8192 | 5 | 1467.6 | 4000 | 2000 | 491,491,... | ... |
| … | … | … | … | … | … | … | … | ... |
| readN | 0 | 8192 | 3 | 1467.6 | 4000 | 3000 | 400,400,... | ... |

**Table 2. Example of a SLOW5 ASCII file with multiple read groups.**

Blue = global header. Yellow = data header. White = data records.

| | | | | | | | | |
|---|---|---|---|---|---|---|---|---|
| #slow5_version | 1.0.0 | | | | | | | |
| #num_read_groups | 3 | | | | | | | |
| @asic_id | 0004A30B00232BEC | | 1004A30B00232BEC | | | 2004A30B00232BEC | | |
| @exp_start_time | 2020-01-01T00:00:00Z | | 2020-01-01T00:00:00Z | | | 2020-01-01T00:00:00Z | | |
| @flow_cell_id | FAH00000 | | FAH00001 | | | FAH00002 | | |
| @run_id | 855cdb | | 855cd1 | | | 855cdc | | |
| … | … | | … | | | … | | |
| #char* | uint32_t | double | double | double | double | uint64_t | int16_t* | ... |
| #read_id | read_group | digitisation | offset | range | sampling_rate | len_raw_signal | raw_signal | ... |
| read-0 | 1 | 8192 | 6 | 1467.6 | 4000 | 4000 | 498,492,... | ... |
| read-1 | 0 | 8192 | 5 | 1467.6 | 4000 | 2000 | 491,491,... | ... |
| … | … | … | … | … | … | … | … | ... |
| read-N | 2 | 8192 | 3 | 1467.6 | 4000 | 3000 | 400,400,... | ... |

#### SLOW5 Header

The SLOW5 header stores metadata regarding the ONT experiment. Header lines start with either '#' or '@'. The header contains two parts: the **global header** (blue fields in tables above) and the **data header** (yellow fields in tables above).

##### *Global header*

The lines starting with '#' form the global header (blue fields above).

The header lines are as follows:

1. The first line of a SLOW5 ASCII file is a key-value pair that specifies the SLOW5 version. The key is separated from the value using a tab '\t'.
2. The second line specifies the number of read groups in the file. Observe that in the single read group file example (**Table 1**), the value for *num_read_groups* is set to 1. In the second example with three read groups (**Table 2**) the value is set to 3.
3. The last line of the header is always the field names for the subsequent per-read records.
4. The second last line of the header specifies the data types of each field for the subsequent per-read records (i.e., for the fields named in the last line of the header). Further information about the fields is provided in the **SLOW5 Data** section below.

##### *Data header*

The header lines that start with '@' form the **data header** (yellow fields above). These header lines contain ONT data attributes that are shared across multiple reads in a sequencing run (read group). For instance, the *run_id* and the *flow_cell_id* are common to all the reads in the read group and are therefore stored in the data header (**Table 1**). These data header lines should always lie after the first two mandatory global header lines and before the last two mandatory global header lines, as illustrated in **Tables 1** & **2**.

For a SLOW5 file containing a single *run_id*, data header lines are key-value pairs delimited by a tab '\t' (**Table 1**). When there are multiple *run_ids* present, the key is followed by a series of values delimited by tabs '\t' (**Table 2**). The first value is for the read group 0, the second value is for the read group 1, the third value is for the read group 2 and so on.

If any attribute value is missing from a given read group a "." is used.

As indicated by the '…' in **Table 1** & **2** after the *run_id* row, many other data header lines may exist, encoding many attributes associated with an ONT sequencing run/s.

The dataset headers are sorted in ascending order based on the native byte values (US-ASCII in C/POSIX locale) of the key. Sorting ensures the SLOW5 file format can easily accommodate the addition or removal of attributes by ONT in the future. A list of possible data header attributes (not an exhaustive list) is provided in **Table 3** below with descriptions based on our understanding of FAST5 files (note: we did not develop FAST5 files, so the definitions here may not be completely accurate).

#### Table 3. Common SLOW5 header attributes.

| Data header attribute key | Description | Example value |
|---|---|---|
| asic_id | Application Specific Integrated Circuit identifier (ASIC) of the flowcel (unique number of the chip), for nanopore tracking. | 213553007 |
| asic_id_eeprom | The identifier of the ASIC's electrically erasable programmable read-only memory (EEPROM) of the flow cell. | 5309577 |
| asic_temp | The temperature in degrees celsius of the ASIC chip, presumably at the start of the sequencing run. | 28.867193 |
| asic_version | The version of ASIC being used. | IA02D |
| auto_update | Indicates whether auto-update in Minknow is enabled or not. | 0 |
| auto_update_source | The link to the Minknow update source. | https://mirror.oxfordnanoportal.com/software/MinKNOW/ |
| barcoding_enabled | Indicates whether barcode demultiplexing is enabled during live basecalling. | 0 |
| bream_is_standard | Bream is one of the software for controlling sequencing. | 0 |
| configuration_version | The MinKNOW core version. | 4.0.13 |
| device_id | The serial ID of the MinION or device position for GridION/PromethION. Device position on GridION/PromethION refers to the ID of the bay (slot where the flow-cell is put) on the device. | X2 |
| device_type | The device type (currently MinION, PromethION or GridION). | gridion |
| distribution_status | Stable vs dev/alpha/beta status. | stable |
| distribution_version | MinKNOW version maybe. | 20.06.9 |
| exp_script_name | These are the scripts used based on what kits are selected in MinKNOW for sequencing. | sequencing/sequencing_MIN106_DNA:FLO-MIN106:SQK-LSK109 |
| exp_script_purpose | Whether run is a real sequencing run or a simulation playback. | sequencing_run |
| exp_start_time | Start time of sequencing run. | 2020-09-08T01:23:21Z |
| experiment_duration_set | Indicates the duration of the experiment selected when starting the sequencing run (presumably in minutes) | 4320 |
| experiment_type | Indicates the type of the experiment, for instance, genomic_dna or rna. | genomic_dna |
| flow_cell_id | Unique ID for the flow-cell, used by ONT to track flow-cell metrics and warranty. | FAN43349 |
| flow_cell_product_code | The type of flow-cell, these will be different based on R9.4.1, R10.3, R9.5, PromethION, etc. | FLO-MIN106 |
| guppy_version | Guppy version being used by MinKNOW. | 4.0.11+f1071ce |
| heatsink_temp | The heat sink that is on the sequencer that is used to control the ASIC temp in degrees celsius most probably at the start of the sequencing run. | 33.996094 |
| hostname | The hostname of the computer doing the sequencing run. | GXB02243 |

| | | |
|---|---|---|
| installation_type | This is the MinKNOW install type. We believe that for GridION and MinION it is nc, and something else for PromethION. | nc |
| local_basecalling | Indicates if live base calling is enabled or not. | 1 |
| local_firmware_file | We currently do not have information on this. | -1 |
| operating_system | The operating system and the version of the computer performing the sequencing run. | ubuntu 16.04 |
| package | We are note sure about this attribute, but it seems to relate to Bream [https://github.com/nanoporetech/minknow_lims_interface]. | bream4 |
| Package_version | We currently do not have information on this. | 6.0.7 |
| protocol_group_id | This is the run name given by the user for the experiment group. | GLFN180082 |
| protocol_run_id | I think this is a unique UUID for the experiment GROUP (just in case the name given by the user is not unique). This is same for each run of the same experimental group. | f2c69573-5fef-43b8-8d81-9cb20634aa7c |
| protocols_version | Not sure exactly what this is, but probably a way MinKNOW tracks it's various protocols for barcoding, kits, etc. | 6.0.7 |
| run_id | The unique run ID for a given run, even in the same experiment group. | 07770780274b0e3703f00d969291b1a37a5a6be1 |
| sample_frequency | Appears to be same as the sampling_frequency in the channel_id group. | 4000 |
| sample_id | Sample ID is the name given by the user for the sample. | NA12878 |
| sequencing_kit | The sequencing kit used, for instance, sqk-lsk109 or sqk-rna002. [https://store.nanoporetech.com/sample-prep.html] | sqk-lsk109 |
| usb_config | Various information about the connection between the flow-cell and the computer. | GridX5_fx3_1.1.3_ONT#MinION_fpga_1.1.1#bulk#Auto |
| version | We currently do not have information on this. | 4.0.3 |

*NOTE:* Many of the attributes in **Table 3** are not used in a typical signal analysis experiment and many are also inconsistent between various FAST5 versions. Although they are unlikely to be used, these attributes are retained by default when converting from FAST5 to SLOW5 format.

#### SLOW5 Data

After the SLOW5 header, the actual data is encoded (white fields in **Tables 1** & **2**, above). Each line contains information about a single read and we refer to this as a record. Each record is made up of several fields that are tab delimited.

As mentioned earlier, the last header line specifies the name of each field. There are two types of fields:

- Primary fields are mandatory and arranged in a strict order. There are 8 primary fields, which are exemplified in **Table 1** & **2** (from the *read_id* field to *raw_signal* field)
- Auxiliary fields are optional and arranged in no strict order. There can be 0 or more auxiliary fields and these are denoted by the '…' after the *raw_signal* field in **Table 1** & **2**.

The second last header line specifies the data type of each primary & auxiliary field. For the primary fields the data types are always the same, whereas the auxiliary field types depend on the fields themselves. The supported data types in SLOW5 are:

- 8-bit, 16-bit, 32-bit and 64-bit signed integers (*int8_t*, *int16_t*, *int32_t*, *int64_t*) and corresponding 1D arrays (*int8_t**, *int16_t**, *int32_t**, *int64_t**)
- 8-bit, 16-bit, 32-bit and 64-bit unsigned integers (*uint8_t, uint16_t, uint32_t, uint64_t*) and corresponding 1D arrays (*uint8_t*, uint16_t*, uint32_t*, uint64_t**)
- IEEE 754 32-bit and 64-bit precision floating point (*float, double*) and corresponding 1D arrays (*float**, *double**)
- ASCII characters (*char*) and ASCII strings (*char*)*. Note: Tabs ('\t' and newline characters '\n' are not allowed in either)

##### *Primary fields*

The 8 primary data fields in SLOW5 format are summarised in **Table 4** below. These fields are mandatory and must be arranged in the order that they appear in **Table 4**.

##### Table 4. Primary data fields in SLOW5 format.

| Field name | Data type | Description | Example value |
|---|---|---|---|
| read_id | char* | A unique identifier for the read. This is a Universally unique identifier (UUID), and should be unique for any read from any device. | 00592138-f120-4ab5-991 6-c5567adb8e29 |
| read_group | uint32_t | Read group identifier. More information in the subsequent text. | 0 |
| digitisation | double | The digitisation is assumed to be the number of quantisation levels in the Analog to Digital Converter (ADC). That is, if the ADC is 12 bit, digitisation is 4096 ($2^{12}$). | 8192.0 |
| offset | double | Assumed to be the ADC offset error. This value is added when converting the signal to pico ampere. | 10.0 |
| range | double | Assumed to be the full scale measurement range in pico amperes. | 1441.389892578125 |
| sampling_rate | double | Sampling frequency of the ADC, i.e., the number of data points collected per second. | 4000 |
| len_raw_signal | uint64_t | The number of samples in the raw signal (length of the raw_signal vector below). | 59676 |
| raw_signal | int16_t* | The raw signal which are the direct acquisition values from the ADC and are comma separated. | 1039,588,588,593,586…. |

Of the 8 primary fields, *read_group* is the only field that does not appear in ONT's FAST5 format but has been introduced in SLOW5. *read_group* identifiers allow reads from multiple sequencing runs to be stored in the same file. *read_group* is essentially an index (0-based index) that specifies where the data header values for a given read are to be found in the data header. For instance, in **Table 2**, *read0* has the *read_group* 1 which means that the second value of the three values for each header attribute contains information for that particular sequencing run (e.g. out of the three values for the *flow_cell_id* key, second one is FAH00001).

In the SLOW5 header, the *num_read_groups* specify how many read groups are present. For instance, in **Table 2**, there are 3 samples in the file and thus *num_read_groups* is equal to 3. Note that the following should always be true: 0 <= *read_group* < *num_read_groups*. *read_group* is always 0 for a single sample file (as it is in **Table 1**).

Datasets are separated into multiple read groups based on the *run_id* (which is a unique string for a sequencing run specified in the data header). The indexing order of the read groups (*read_group)* is determined by the order the FAST5 files are parsed during FAST5 to SLOW5 conversion. This *read_group is* an internal index used for enumerating. This index allows more efficient enumeration (less computation and saves disk space) than performing string comparisons if *run_id* string was stored in the data record for every read instead.

Primary fields contain all the information required for a typical nanopore signal-level analysis. The raw signal can be easily converted to pico-ampere using the following equation:

```
signal_in_pico_ampere = (raw_signal + offset) * range / digitisation
```

##### *Auxiliary fields*

SLOW5 files may contain 0 or more auxiliary data fields, some common examples of which are provided in **Table 5** below. These fields are optional and not bound by any strict order.

**Table 5. Common auxiliary data fields in SLOW5 format.**

| Field name | Data type | Description | Example value |
|---|---|---|---|
| channel_number | char* | The channel number. A flow cell has multiple channels allowing multiple DNA/RNA strands to be sequenced in parallel. For instance, a MinION flow cell has 512 channels and thus can sequence 512 strands in parallel. | 504 |
| median_before | double | The measure of the median current level for the data preceding the read. In most cases this can be used as an estimate of the open pore level. The open-pore state is when there is no strand inside the pore. | 238.78225708007812 |
| read_number | int32_t | This number is unique within the channel. | 17981 |
| start_mux | uint8_t | The MUX setting for the channel when the read began. Each channel contains one or more wells. For instance, a MinION flow cell has 4 wells per channel. The wells within a channel are connected to a multiplexer (MUX), a switch that controls which of the four wells in the channel is controlled and read out for sequencing. | 4 |
| start_time | uint64_t | The start time of the read. The unit for *start_time* is 'number of sampling events', so *start_time* has to be divided by sampling rate (sampling_rate) to get the start time in seconds (i.e. the time since the run was started) | 335845487 |

Auxiliary fields contain all per-read information from ONT FAST5 files that we do not consider primary data fields (i.e., attributes that are not commonly used in signal-level analysis). If a value for a particular auxiliary field is unavailable for a given read it is represented with a ".".

It is important to note that auxiliary fields can be in any order, meaning the user should not rely on their order and instead should enumerate based on the field names and data types specified in the header.

Any future per-read attributes added to FAST5 by ONT will be included as auxiliary fields in SLOW5. If ONT drops any attribute from FAST5, it will also be dropped in SLOW5.

The auxiliary fields are separated from each other and from the primary fields by using a tab '\t' as a delimiter. The elements in a field of 1D array data type (except *char** strings) are delimited by commas. Strings are stored as a series of characters, as usual, and the null terminating character is not stored.

### BLOW5

A SLOW5 binary file or a BLOW5 file is the binary counterpart to a SLOW5 ASCII file. The file extension is *.blow5*. In BLOW5 format all multi-byte numbers are stored in little-endian, regardless of the machine's endianness.

A BLOW5 file can be either uncompressed or compressed. BLOW5 implementation currently supports *zlib (DEFLATE* algorithm*)* as the compression method, but the format itself is designed such that any other compression method can be seamlessly integrated. The BLOW5 header specifies whether the file is uncompressed or compressed, and the compression method that is applicable.

#### BLOW5 Header

The fields of the BLOW5 header are displayed in **Table 6** below. Note that despite being shown in a table for clarity, the fields in a BLOW5 file are stored serially in the exact order as they are in **Table 6**, without any tabs or newlines to separate the fields. The byte offset in the file (first column) and the size of the field in bytes (second column) are used to locate a particular field within a BLOW5 file.

The first field, the magic number, is a 6 byte string `"BLOW5\1"` used as a signature to identify the file format. The next three fields are for storing the BLOW5 file version and the value here is the same as in the SLOW5 ASCII counterpart. The 5th field indicates if the BLOW5 file is compressed or not and the compression method used if it is compressed. The 6$^{th}$ field is the number of read groups in the file, which have the same value and meaning as in the SLOW5 ASCII counterpart (described above). Finally, 50 bytes are reserved for future fields.

Offset 64 contains an integer field that indicates the size of the upcoming variable-sized field, the SLOW5 ASCII header. The next field is the SLOW5 ASCII header, which is the same as in a SLOW5 ASCII file, with the following exceptions:

1. The first line of the SLOW5 header specifying the `#slow5_version` is removed as this is already stored at the beginning of the BLOW5 header;
2. The second line of the SLOW5 header specifying the `#num_read_groups` is also removed as this is also stored at the beginning of the BLOW5 header;;

Apart from these exceptions, the complete SLOW5 ASCII header is stored, including tabs, newlines and the starting characters “#” and “@”. Note that all header data values will be converted to ASCII strings, despite the data type for the corresponding fields in FAST5 files.

**Table 6. Structure of a BLOW5 file header.**

| Offset | size (bytes) | Description | data type | Value |
|---|---|---|---|---|
| 00 | 6 | Magic number | char[6] | "BLOW5\1" |
| 06 | 1 | Major version number | uint8_t | 0 to 255 |
| 07 | 1 | Minor version number | uint8_t | 0 to 255 |
| 08 | 1 | Patch version number | uint8_t | 0 to 255 |
| 09 | 1 | Compression method | uint8_t | 0 to 255<br>(0 for uncompressed binary, 1 for zlib) |
| 10 | 4 | Number of read groups | uint32_t | 0 to $2^{32}$-1 |
| 14 | 50 | Reserved for future | - | - |
| 64 | 4 | size of the SLOW5 header (without null character) | uint32_t | 0 to $2^{32}$-1 |
| 68 | | The plain text header of the SLOW5 (null character not stored; #slow5_version and #num_read_groups are removed as they are already in the binary header) | char[] | |

#### BLOW5 Data

The SLOW5 data records are serially stored in binary format with each record individually compressed using the compression method specified in the header (not compressed if it is not specified in the header). Note that each record is individually compressed to allow efficient parallel access to different records simultaneously.

**IMPORTANT:** Each BLOW5 record is preceded by the size of the upcoming BLOW5 record in bytes (the size of the compressed record if compressed), which is an 8-byte uint64_t type unsigned integer. Storing this size is useful for faster and easier indexing of a BLOW5. We will refer to this special field as "*len_blow5_rec*" from here onwards.

The fields in an uncompressed BLOW5 record are displayed in **Table 7** below.

**Table 7. Structure of a BLOW5 record.**

| Size (bytes) | Description | Data type |
|---|---|---|
| 2 | string length of the read ID (without null character) | uint16_t |
| <variable> - based on preceding value | read ID (null character not stored) | char* |
| 4 | read group | uint32_t |
| 8 | digitisation | double |
| 8 | offset | double |
| 8 | range | double |
| 8 | sampling_rate | double |
| 8 | len_raw_signal | uint64_t |
| <variable> - based on preceding value | raw_signal | int16_t* |
| | <auxiliary fields> | |

The first field is a uint16_t integer that specifies the size of the upcoming *read_id* string. Then comes the eight primary data fields explained under the SLOW5 ASCII section (see above), but now stored in binary. Note that the *raw_signal* field, which was a comma separated list in SLOW5 ASCII, is now a series of int16_t integers (each 2 bytes in size) stored serially without commas. The size of the *raw_signal* field in bytes in **Table 7** is determined by the product of the *len_raw_signal* and the size of int16_t, which is 2.

The *raw_signal* field in a BLOW5 record is followed by the auxiliary fields, as described above. The fields are stored in the same order and datatypes as specified in the header.

Primitive data types (*int8_t*, uint8_t, *int16_t*, u*int16_t*, *int32_t*, uint32_t, *int64_t, uint64_t, float, double*, *char*) are stored such that: *int8_t*, uint8_t and char taking 1 byte; *int16_t* and u*int16_t* taking 2 bytes, *int32_t*, uint32_t and *float* taking 4 bytes; and, *int64_t, uint64_t and double* taking 8 bytes as shown in **Table 8** below. Any missing data field (represented by a '.' in SLOW5 ASCII) is represented in BLOW5 by using the value stated in column 3 in **Table 8**. This special value that represents a missing value cannot be used to represent the real value.

Auxiliary fields of 1D array data types are stored with the length of the 1D array (the number of elements in the 1D array, not the size in bytes) in the form of an 8 byte unsigned integer (uint64_t) preceding the actual data in the array. The elements in 1D arrays are stored sequentially without any delimiting commas. The size of the array field in bytes is determined by the product of the length of the 1D array and the size of the corresponding primitive data type. A missing array field (“.” in SLOW5 ASCII) is represented by storing 0 as the length of the array.

**Table 8. Primitive data types used in BLOW5 format.**

| **Data type** | **size (bytes)** | **Missing value representation** |
|---|---|---|
| *int8_t* | 1 | INT8_MAX = 2^7-1 |
| int16_t | 2 | INT16_MAX = 2^15-1 |
| *int32_t* | 4 | INT32_MAX = 2^31-1 |
| *int64_t* | 8 | INT64_MAX = 2^63-1 |
| uint8_t | 1 | UINT8_MAX = 2^8-1 |
| uint16_t | 2 | UINT16_MAX = 2^16-1 |
| uint32_t | 4 | UINT32_MAX = 2^32-1 |
| uint64_t | 8 | UINT64_MAX = 2^64-1 |
| *float* | 4 | generic NaN value returned by *nanf(“”)* |
| *double* | 8 | generic NaN value returned by *nan(“”)* |
| *char* | 1 | '\0' |

#### BLOW5 Footer

A BLOW file should always end with the end of file (EOF) marker “5WOLB”. This is useful for detecting file truncation.

### SLOW5 INDEX

A SLOW5 index is a binary file that contains an index to facilitate random access to a SLOW5 ASCII or BLOW5 file based on the *read_id*. The extension of an index for a SLOW5 ASCII file is *.slow5.idx* and for a BLOW5 file is *.blow5.idx*. A SLOW5 index always takes the same binary form as described below, irrespective of whether it is for a SLOW5 ASCII or BLOW5 file.

#### SLOW5 Index Header

##### Table 9. SLOW5 index header structure.

| Offset | size (bytes) | Description | data type | Value |
|---|---|---|---|---|
| 00 | 9 | Magic number | char[9] | "SLOW5IDX\1" |
| 09 | 1 | Major version number | uint8_t | 0-255 |
| 10 | 1 | Minor version number | uint8_t | 0-255 |
| 11 | 1 | Patch version number | uint8_t | 0-255 |
| 12 | 52 | Reserved for future | - | - |

Note: The SLOW5 index version is the same as that of the SLOW5 version in the corresponding SLOW5 file.

#### SLOW5 Index Data

The SLOW5 index data records start from offset 64 of the file. The index should have a single record for every record in the corresponding SLOW5/BLOW5 file. An individual SLOW5 index record takes the following form:

##### Table 10. SLOW5 index data structure.

| Size (bytes) | Description | Data type |
|---|---|---|
| 2 | String length of the read ID (without null character) | uint16_t |
| <variable> - based on preceding value | Read ID (null character not stored) | char* |
| 8 | For ASCII SLOW5: byte offset in the SLOW5 ASCII file that corresponds to the beginning of the data record.<br><br>For BLOW5: byte offset in the BLOW5 file that corresponds to the beginning of the *len_blow5_rec* that precedes the data record. | uint64_t |
| 8 | For ASCII SLOW5: size of the SLOW5 ASCII data record in bytes.<br><br>For BLOW5: size of the BLOW5 data record in bytes (the size of the compressed record if compressed) + the size of the *len_blow5_rec* preceding the record (which is 8 as the datatype of *len_blow5_rec* is uint64_t). | uint64_t |

#### SLOW5 Index Footer

A SLOW5 index file should always end with the end of file marker *"XDI5WOLS"*. This is useful in detecting file truncation.

### RATIONALE BEHIND SLOW5 DESIGN DECISIONS

In this section we provide the rationale behind certain design decisions and why these were preferred over other potential solutions. Please note that some of the following discussions are pretty technical and not for the faint-hearted.

- Why does SLOW5 have two types of fields, primary and auxiliary?
  - Primary fields are the essential elements of signal-based analysis. These essential elements are provided as primary fields for easy and quick accessibility.
  - Auxiliary fields are very application-specific and not generally used in existing signal-based analyses. Keeping these fields separate prevents convoluting the primary fields. Also, auxiliary fields can be in any order and are optional. Therefore, the SLOW5 format will not break when ONT adds or removes a field, and users can optionally choose to discard the auxiliary fields during FAST5 to SLOW5 conversion, in order to reduce file size and complexity.

- Why is SLOW5 one big file opposed to a number of small files?
  - Modern file systems support bigger files. With disk sizes continually growing, this will be increasingly true in the future.
  - Random accesses would require repeated expensive file open and close operations if multiple files are used (the default number of maximally open files in Linux is typically 1024).
  - In the case of a user requiring to perform process-level parallelism on a per-file basis, they could use *slow5tools split* to quickly split the files.
  - When archiving, users tend to tar the files into a single ball anyway if there were multiple files. So why not just create a single file that can be directly archived?
  - Most bioinformatics users are familiar with working on a single large file for a given sample in FASTQ, BAM or VCF format, so we thought it would be good to follow this approach.

- Why does SLOW5 support multiple sequencing runs in the same file?
  - In nanopore sequencing experiments, it is quite common to run more than one flow cell on a given sample, or create a new run id when a flow cell is washed and reloaded during a run. Allowing data from multiple *run_ids* to be stored in a single SLOW5 file means developers do not have to deal with manually accepting multiple files when analysing data from more than one run. It is generally more convenient to have all the data in a single file.

- Why are empty fields in SLOW5 ASCII represented by "."?
  - SLOW5 ASCII is only for human readability and having a "." is easier to read than empty fields. This is also easier to parse when using tools like *awk, sed* and *cut*. Popular formats like SAM, BED and VCF use "." for empty fields, so we chose to stick to this convention.

    - If a single "." is to become a valid field value (unlikely) in FAST5 which is not the case at the moment, we would introduce a workaround such as using "\." or ".." in the future.

- Why is there a version number for the slow5 index?
    - To make it future proof.
    - In the future, we can provide alternate *btree* based indexing for memory efficiency if required.

- Why does BLOW5 use little endian storage?
    - All modern systems seem to use little endian. We are not aware of any big endian systems still in use.

- Why is a tab used as the delimiter in SLOW5 ASCII?
    - Tab separators are easy to parse with *awk, sed, cut* and other command line tools. This also mimics the convention used in SAM, VCF, BED and other genomics formats.

- Why are # and @ used in the SLOW5 header?
    - To distinguish the SLOW5 format related attributes (SLOW5 global header) from nanopore related attributes (SLOW5 data header) we use the two symbols # and @. Both of those characters are not supposed to be used in read identifiers and therefore there is no confusion with the data records.
    - We considered '##' for SLOW5 global header and a '#' for the SLOW5 data header but we decided against this because if ONT introduces an attribute name that starts with a '#', this would cause a lot of problems for SLOW5.

- Why is native byte order sort used for the attribute names in the SLOW5 data header?
    - There are many different data attributes and these are quite hard to keep track of because they differ between different FAST5 versions. Sorting these makes it easy for a human to quickly locate the information they are after.
    - To prevent adhoc ordering which would make it difficult to parse.

- Why doesn't SLOW5 support the analysis group in FAST5 files?
    - SLOW5 is meant for storing raw signal data. Storing analysis data (e.g., basecalled FASTQ records) would convolute the file format. We believe those post processing information should be a separate file, as is the case in other areas of bioinformatics where, for example, raw reads (FASTQ), alignments (BAM) and variants (VCF) are stored in separate files with specific formats.

- Why is a SLOW5 index always in binary and no ASCII version?
    - For fast loading and space saving.
    - The index is primarily read by a computer and not particularly useful to a human.

- Do any of the float/double fields in SLOW5 ASCII become lossy when they are converted to ASCII strings?

    - Yes, some of them do (for example recurrent decimals). However, this is not an issue when FAST5 is directly converted to BLOW5 as floats/doubles are directly stored in binary. SLOW5 ASCII is meant for viewing the binary counterpart BLOW5 by humans and not meant to be used for data archiving or processing. We recommend using the default conversion setup in *slow5tools f2s* that converts FAST5 files to zlib compressed BLOW5 files initially and the later use *slow5tools view*.
    - In ASCII we could have stored the float/double fields in hexadecimal to make lossless, but then this is not as readable to humans as a natural representation like x.y

- Do the values stored in the data header attributes become lossy as floats are also converted to string?
    - Currently all the data header attributes in SLOW5 are stored as ASCII strings in FAST5 as well. So at the moment there is no loss.

- What happened to the duration attribute in FAST5?
    - The duration attribute has a bad history. A few years ago this used to be the length of the signal in seconds and now this is used by ONT to represent the length of the signal in terms of number of samples. To avoid ambiguity we store the length of the signal in terms of the number of samples in the field *len_raw_signal* in SLOW5, which is equivalent to the value in the duration attribute in FAST5.
    - If ONT decides to make the duration in seconds again, we can add this as an auxiliary field for SLOW5 while keeping the *len_raw_signal* intact in SLOW5 as the length of the signal is essential in signal analysis.

- What if the end of file markers “5WLOB” or “*XDI5WOLS*” occur in the middle of a file?
    - This is possible to happen if the data by any chance matches this pattern in binary. However, this is not an issue as the end of the file marker in BLOW5 is used to detect file truncation. That is, we check if the end of the file marker is present only if the EOF has been reached.
    - The case that the data at the end of a truncation is translated to an end of marker is extremely rare.

- Why are certain fields such as “digitisation” that seem to be identical across all reads in a given sequencing run present in data records, opposed to being header data attributes?
    - These are essential values for converting the raw signal. So it is quite convenient to have them adjacent to the raw signal. Also in case something happens to the header, the records will still be usable.
    - In the future, this digitisation attribute may no longer be the same across reads (as ONT stores this redundantly for each read unlike the header data attributes which are symbolic links).

- Are options header lines that start with ‘#’ supported in SLOW5?
    - No. Optional lines would complicate parsing and can include complicated situations where different users starting to use a header of their own and later causing confusions. If the requirement comes, we will introduce this in a future version.

- What if the forbidden ‘\t’ and ‘\n’ in data header values and auxiliary fields ever should become a valid character in FAST5?
    - At the moment they are forbidden to keep the file format simple. If this ever happens, in future versions we will allow ‘\\t’ or something.

### MISCELLANEOUS

#### SLOW5 versioning

While forward-compatibility cannot be ensured, backward compatibility will be maintained where possible.

Versioning follows the *major.minor.patch* approach.

- The *patch* version is incremented for backwards compatible bug fixes.
- The *minor* version is incremented for backward compatible newer features and functions.
- The *major* version is incremented when potentially backward incompatible changes are introduced.

There are two independent tracked versions:

1. slow5 file and slow5 index file version
2. *slow5lib*, *pyslow5* and *slow5tools* versions

The slow5 file and slow5 index file version is independent from the *slow5lib*, *pyslow5* and *slow5tools* version and is used for checking compatibility.

*slow5lib, pyslow5 and slow5tools* versions are independently patched while maintaining compatibility, and are version synced during any stable release.
